# The utility of mortality hazard rates in population analyses

**DOI:** 10.1101/216739

**Authors:** Torbjørn Ergon, Ørnulf Borgan, Chloé Rebecca Nater, Yngvild Vindenes

**Author notes:** Corresponding author, /.

## Abstract

1. Mortality is a key process in ecology and evolution, and much effort is spent on development and application of statistical and theoretical models involving mortality. Mortality takes place in continuous time, and a fundamental representation of mortality risks is the mortality hazard rate, which is the intensity of deadly events that an individual is exposed to at any point in time. In discrete-time population models, however, the mortality process is represented by survival or mortality probabilities, which are aggregate functions of the mortality hazard rates within given intervals. In this commentary, we argue that focussing on mortality hazard rates, also when using discrete-time models, aids the construction of biologically reasonable models and improves ecological inference.
2. We discuss three topics in population ecology where hazard rates can be particularly useful for biological inference, but are nevertheless often not used: (i) modelling of covariate effects, (ii) modelling of multiple sources of mortality and competing risks, and (iii) elasticity analyses of population growth rate with respect to demographic parameters. To facilitate estimation of cause-specific mortality hazard rates, we provide extensions to the R package ‘marked’.
3. Using mortality hazard rates sometimes makes it easier to formulate biologically reasonable models with more directly interpretable parameterizations and more explicit assumptions. In particular, interpretations about relative differences between mortality hazard rates, or effects of relative changes in mortality hazard rates on population growth (elasticities), are often more meaningful than interpretations involving relative differences in survival (or mortality) probabilities or odds.
4. The concept of hazard rates is essential for understanding ecological and evolutionary processes and we give an intuitive explanation for this, using several examples. We provide some practical guidelines and suggestions for further methods developments.

## Introduction

Mortality takes place in continuous time, but statistical and theoretical models in ecology are often formulated on a discrete time-scale, where the mortality process is represented by **survival** or **mortality probabilities** (terms in bold at first occurrence are defined in the Glossary). While **mortality hazard rate** (*h*(*t*)) is the instantaneous intensity of deadly events that an individual is exposed to (Box 1), the probability of surviving a given interval from time *t*_1_ to time *t*_2_ (*S*_*t*_1_→*t*_2__) is a function of the integrated hazard rate over the interval,

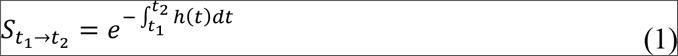

(Box 1). The integral in this aggregate function can be replaced by the average hazard rate within the interval multiplied by the length of the interval such that survival probability in interval *j* becomes

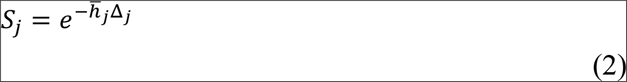

where Δ_*j*_ is the length of the time-interval (usually one time-unit) and the unit of measurement for the time-averaged hazard rate, *h̅*_*j*_, is number of deadly events per time-unit (Collett 2014). Such interval-specific survival probabilities and time-averaged hazard rates are often sufficient for specific analyses where fluctuations within intervals (e.g. due to seasonal or daily cycles) are not relevant.

Although survival probabilities are simple transformations of time-averaged mortality hazard rates (eq. (2)), there are some important differences in how these two measurements can be interpreted and modelled. The survival probability is a unit-free measurement on an **absolute scale**, meaning that no transformation can be performed without changing its meaning (Houle *et al.* 2011). For example, if *S* is the probability of surviving a whole year, then *S*^1/2^ can be interpreted as the probability of surviving half the year, assuming that survival probabilities in the first and second half of the year are equal. The mortality hazard rate, on the other hand, is a **ratio-scaled measurement**, and its unit of measurement may thus be changed (e.g., from ‘year^−1^’ to ‘month^−1^’) by simply multiplying with a positive number (just like a given distance can be measured in ‘meters’ or ‘feet’) (Houle *et al.* 2011).

Any model based on survival probabilities can be rewritten in terms of time-averaged hazard rates using eq. (2). The relationship between cause-specific mortality probabilities and hazard rates is somewhat more complex, and we will return to this later. In the fields of statistical modelling of capture-recapture data and structured population modelling, there is a tradition for basing inference on survival probabilities, odds-ratios and effects of relative changes (elasticities) in survival probabilities (Caswell 2001; McCrea & Morgan 2015). In other fields, such as fisheries, medicine and human demography, it is more common to model and interpret mortality hazard rates and hazard ratios (Quinn & Deriso 1999; Keyfitz & Caswell 2005; Collett 2014). Mortality hazard rates are also often used in life-history theory (e.g. Charnov 1993). Sometimes, using mortality hazard rates makes it easier to formulate biologically reasonable models with more directly interpretable parameterizations and more explicit assumptions, and our aim is to illustrate this. Our philosophy is that models should not just fit the data well (often many models fit the available data equally well), but also have a well-argued theoretical foundation and support meaningful and clear biological interpretation. Although a substantial lack of fit to the data should prompt a reconsideration of the model (Mac Nally *et al.* 2017), it is not given that the model with the best fit (in an absolute or relative sense) should be preferred. Hence, in the following, we will only discuss ‘a priori’ modelling choices.

In the first section below, we argue that log-hazard (loglog-link) models are generally more suited to describe effects of covariates on survival probabilities than the commonly used log-odds (logit-link) models. Under the second heading, we discuss and illustrate the use of hazard rate models to study multiple competing sources of mortality. In the third section, we discuss the utility of mortality hazard rates in elasticity analyses of demographic models. Finally, we conclude with some perspectives on future developments. All sections are accompanied by further details, including R-code, in the online Supporting Information.

### Log-linear models of mortality hazard rates vs. logit-linear models of survival probabilities

Understanding effects of different covariates (e.g. relating to climate or individual properties) on survival/mortality is a central aim in ecology and evolution. Such effects are typically estimated using linear models of survival probabilities transformed by a link-function (Lebreton *et al.* 1992). When modelling effects of covariates on time-averaged hazard rates (eq. 2), **log-linear models** are attractive because effects then represent relative differences (i.e., **hazard ratios**) which are invariant to the time scale used and have a clear interpretation (Box 1). A log-linear model for time-averaged mortality hazard rates is equivalent to using a **loglog-link** on survival probabilities (Box 1). In comparison, when survival probabilities are modelled by the more commonly used **logit-link** function, the exponential of **linear contrasts** may be interpreted as **odds-ratios** (Box 1 and Appendix S1 in the Supporting Information).

While most practitioners are presumably aware of the different functional forms implied by the choice of link-function when using continuous covariates, there seems to be less awareness of the fact that parameterising the models with survival probabilities representing different durations (e.g. months instead of years) also changes the functional forms when using any other link-function than the loglog-link (Figure 1; Appendix S1 includes R code to reproduce all figures in the article and provides additional explanations). That is, when *not* using a loglog-link for survival probabilities, selecting the time-unit for the analysis (which can be set in commonly used computer programs such as MARK (White & Burnham 1999)) is part of the model definition. In many studies, it is natural to model survival on an annual time scale as this represents survival over a full seasonal cycle. However, for species with a life-expectancy of much less than a year, such as many small rodents, yearly survival is not meaningful, and one may choose, somewhat arbitrarily, to model weekly or monthly survival instead. If a logit-link is used in such cases, one will get different results depending on the choice of interval length that the survival probabilities in the model represent. Using a loglog-link for survival, which implies a log-linear model for the time-averaged mortality hazard rate (Box 1), avoids the problem of selecting the time-spans that the survival probabilities represent because survival over any given interval is then invariant to the interval length used in the modelling of covariate effects (Figure 1; Appendix S1). Further, mortality hazard rates provide a cohesive basis for fitting models of senescence (Gaillard *et al.* 2004), to model seasonal changes in mortality as continuous functions of time (Ergon, Yoccoz & Nichols 2009), to allow transitions between individual states in continuous time (Miller & Andersen 2008; Ergon, Yoccoz & Nichols 2009; Conn, Cooch & Caley 2012; Choquet *et al.* 2017), or to accommodate analysis of data where individuals are not encountered during discrete occasions (Choquet *et al.* 2017).

**Figure 1.**
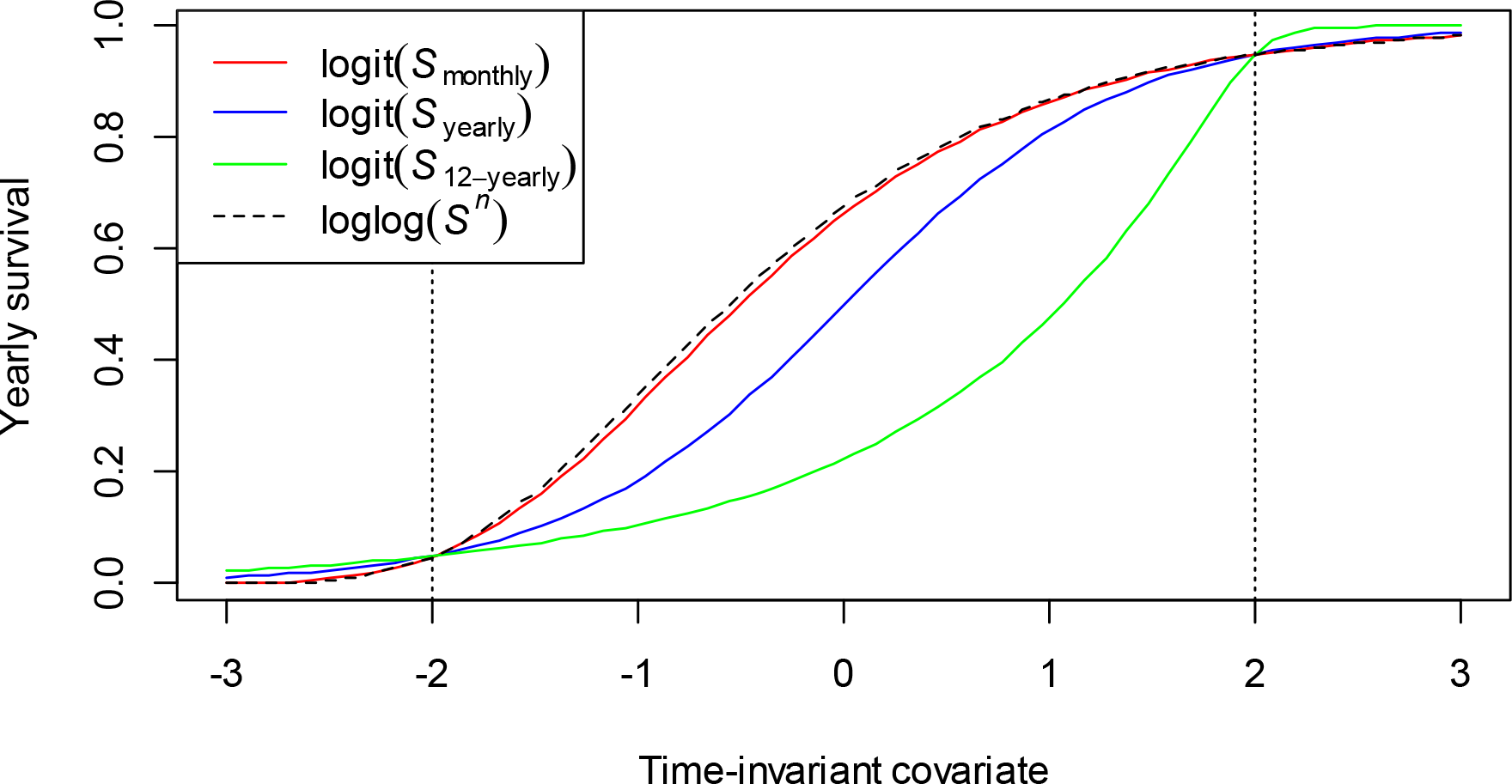
Functional shape of the relationship between yearly survival and a time-invariant covariate (e.g. individual specific) when using the logit link-function on survival probability over different time interval lengths (solid coloured lines; see legend). When the logit-link is applied to monthly survival (red line), it is assumed that probability of survival is the same in all of the 12 months of the year, such that 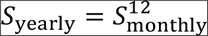. Similarly, when the logit-link is applied to survival over 12 years, 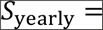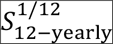 (green line). To facilitate comparison of functional shapes, slope and intercept parameters are defined such that yearly survival probability is 0.05 at covariate value −2 and 0.95 at covariate value 2. The dashed line shows the loglog-linear relationship between yearly survival and the covariate, which is invariant to the interval length for the survival probability. When the logit-linear relationship applies to survival probabilities over short intervals, it approaches the loglog-linear functional form (see red line).

While survival probabilities, as well as their absolute and relative differences, may provide meaningful inference as long as one acknowledges that these measurements depend on the defined interval lengths, differences in survival expressed as odds-ratios are difficult to interpret in a biological context. For example, consider an environment where yearly juvenile survival is 0.25 (odds = 0.25/(1-0.25) = 1/3) and yearly adult survival is 0.50 (odds = 0.5/(1-0.5) = 1/1). Here the odds of surviving one year is 3 times higher for adults compared to juveniles, but the odds of surviving two years is 5 times higher for adults than juveniles (1/3 vs. 1/15). Now consider an improved environment where yearly survival of juveniles and adults are respectively 0.50 and 0.71, the odds ratios for surviving one year is 2.4 (and 3.0 for surviving two years), suggesting a smaller difference between juveniles and adults than in the previous environment. In both environments, however, the time-averaged mortality hazard rate (eq. 2) for juveniles is 2 times higher than the mortality hazard rate for adults, meaning that the intensity at which deadly events occur is twice as high for juveniles compared to adults, irrespective of the environment and the time-scale used. Hence, as illustrated in Figure 2, an odds-ratio for any given hazard ratio does not only depend on the length of the time-interval that the odds apply to, but also on the survival probability of the reference group. In contrast, hazard ratios have a clear biological interpretation, and facilitate meaningful comparisons of effect sizes (Box 1).

**Figure 2.**
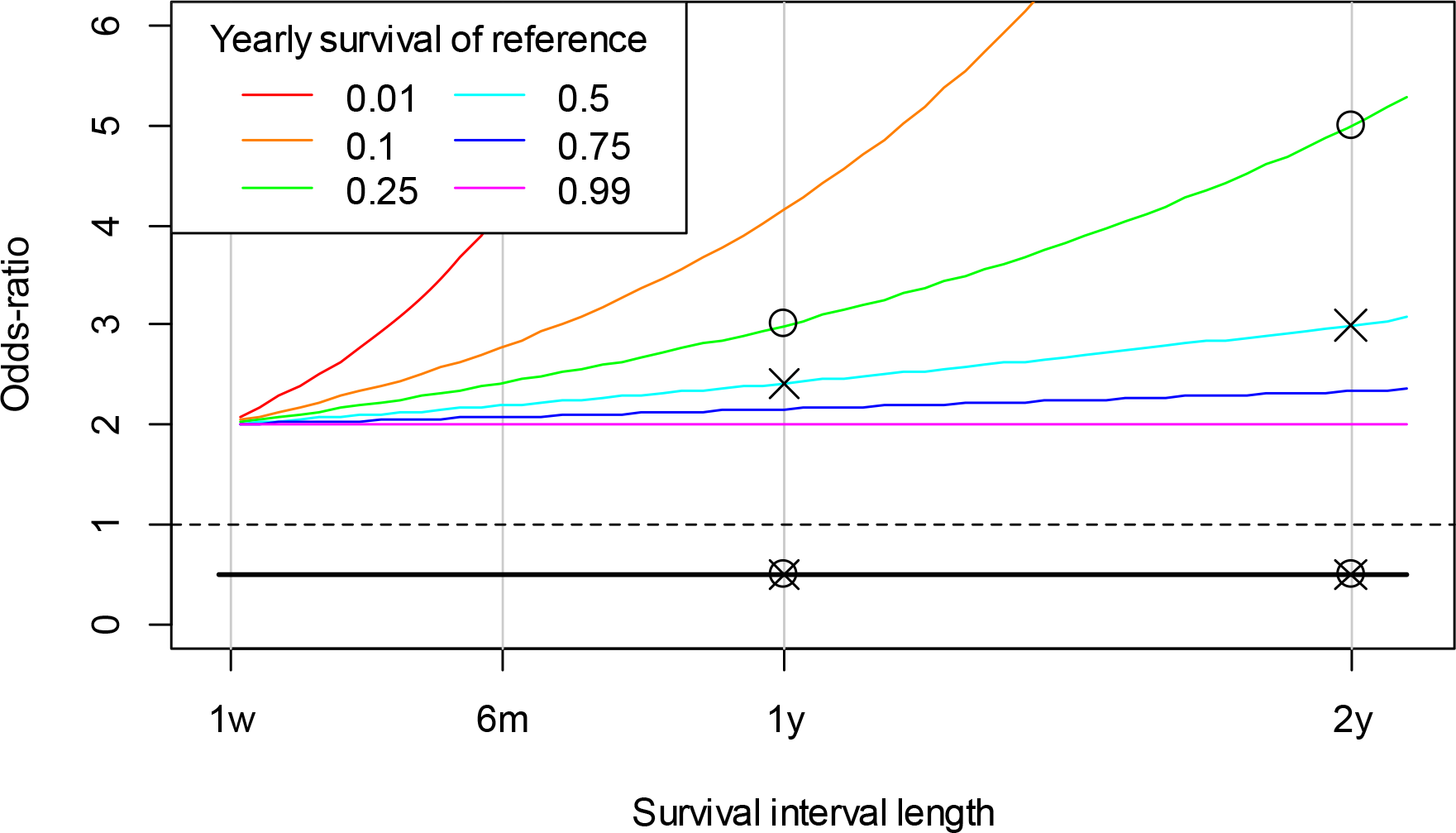
Contrasts between two survival probabilities expressed as odds-ratios as a function of time interval length for survival (x-axis), when the mortality hazard ratio is 0.5 (indicated with a thick bla line) and yearly survival probability of the reference varies according to the colour code in the legend Plotted points refer to the example in the text (circles represent the original environment, while cross represent the improved environment). Vertical lines are plotted at 1 week, 6 months, 1 year and 2 years.

### Modelling of multiple sources of mortality and competing risks

Discerning between different sources of mortality is a key challenge in ecology, evolutionary biology and population management (Schaub & Lebreton 2004; Péron 2013; Wolfe *et al.* 2015). To understand selective forces and population dynamics, it is also important to examine how different sources of mortality co-vary and interact (Nichols *et al.* 1984; Boyce, Sinclair & White 1999; Koons *et al.* 2014). For example, an essential question is whether increased hunting or harvesting will lead to a reduction (or potentially an increase) in mortality due to other causes. Such effects can result from density dependent responses or non-random removal of weaker (or stronger) individuals (i.e., selection) in a heterogeneous population. Changes in one source of mortality may also lead to evolutionary changes in other mortality rates or life-history traits. For example, age-dependent mortality patterns affect optimal investments in development and maintenance/longevity as well as reproductive effort at different ages (Charlesworth 1980; Caswell 2007a; Koons *et al.* 2014).

When studying multiple sources of mortality, it is essential to recognize that if the hazard rate representing one cause of mortality (say, hunting) increases, the probability of dying from any other cause will automatically be reduced, even if the mortality hazard rates representing the other causes remain unchanged. This is simply because it has now become more likely that the individual will die of this cause (hunting) before other deadly events can occur (Appendix S2 in the Supporting Information). We may say that the different causes of mortality “compete” over killing the individual first, and the probability of “winning” depends on how “strong” the competitors are (referred to as “competing risks” in survival analysis (e.g. Collett 2014)).

If we assume that there are *K* causes of mortality and the mortality hazard rates for each cause *k*, *h*_*k*_(*t*), remain proportional within a time interval, such that the fraction 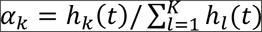 remains constant for all causes throughout the interval, it can be shown (see Appendix S2) that the probability of dying from cause *k* is

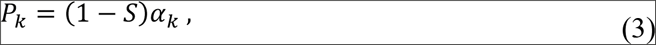

where *S* is the probability of surviving the interval (from *t_1_* to *t_2_*) defined to have length 1. That is,

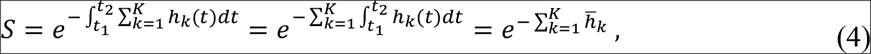

where *h̅*_*k*_ is the time-averaged hazard rate for cause *k* (eq. (2)). In the most general model these parameters would be interval- and individual-specific, but indices for interval and individual are omitted here for simplicity. Note that when the hazard rates *h*_*k*_(*t*) remain proportional within an interval, the relative differences between them are the same as the relative differences between the time-averaged hazard rates. Hence, we can also interpret *α*_*k*_ as

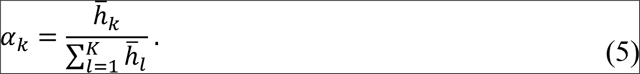

When information about cause of death (e.g., harvesting) is available for at least a subset of marked individuals in the population, multi-state capture-recapture models can be fitted to the data to address questions relating to e.g. compensatory effects of harvesting or age-related patterns in mortality from different causes (Servanty *et al.* 2010; Koons *et al.* 2014). Such models involve probabilities of remaining in an “alive-state” (survival probability *S*) or transitioning to any of the absorbing death-by-cause states (probabilities *P*_*k*_’s). From equations (3), (4) and (5) we see that there are at least three distinct ways to parameterize these transition probabilities when hazard rates within intervals are assumed to be proportional: [1] with *S* and *P*_*k*_’s under the constraint that 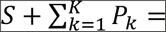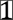[2] with *S* and *α*_*k*_’*s* under the constraint that 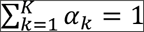; or [3] with the cause-specific time-averaged mortality hazard rates *h̅*_*k*_’s. The relationships between these three parameterizations are given in Table 1. Note that when constraining these individual- and interval-specific parameters by functions of individual age or covariates, these alternative model formulations will generally lead to different models.

**Table 1.**
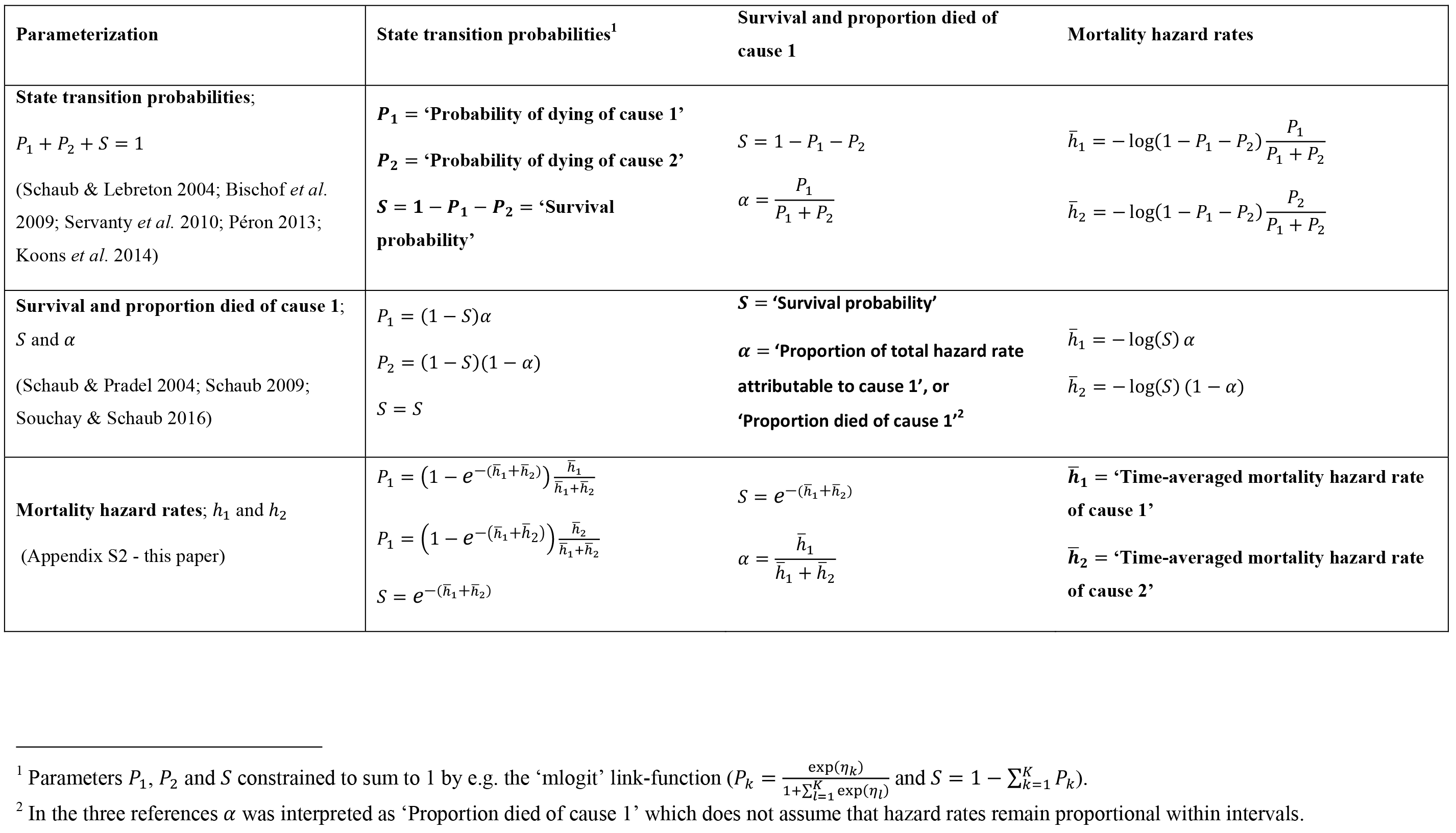
Matrix of three alternative parameterizations of state transition probabilities in a multi-state model of two mutually exclusive and collectively exhaustive causes of mortality, assuming that cause specific hazard rates remain proportional within intervals of length 1. Definitions of the parameters are given in the diagonal cells. These parameters may be interval and individual specific, and may be modelled as random effects or as functions of covariates in a hierarchical model.

| Parameterization | State transition probabilities<sup>1</sup> | Survival and proportion died of cause 1 | Mortality hazard rates |
| --- | --- | --- | --- |
| **State transition probabilities;**<br>$P_1 + P_2 + S = 1$<br>(Schaub & Lebreton 2004; Bischof *et al.* 2009; Servanty *et al.* 2010; Péron 2013; Koons *et al.* 2014) | $P_1 = \text{'Probability of dying of cause 1'}$<br>$P_2 = \text{'Probability of dying of cause 2'}$<br>$S = 1 - P_1 - P_2 = \text{'Survival probability'}$ | $S = 1 - P_1 - P_2$<br>$\alpha = \frac{P_1}{P_1 + P_2}$ | $\bar{h}_1 = -\log(1 - P_1 - P_2) \frac{P_1}{P_1 + P_2}$<br>$\bar{h}_2 = -\log(1 - P_1 - P_2) \frac{P_2}{P_1 + P_2}$ |
| **Survival and proportion died of cause 1;**<br>$S$ and $\alpha$<br>(Schaub & Pradel 2004; Schaub 2009; Souchay & Schaub 2016) | $P_1 = (1 - S)\alpha$<br>$P_2 = (1 - S)(1 - \alpha)$<br>$S = S$ | $S = \text{'Survival probability'}$<br>$\alpha = \text{'Proportion of total hazard rate attributable to cause 1', or 'Proportion died of cause 1'}^2$ | $\bar{h}_1 = -\log(S) \alpha$<br>$\bar{h}_2 = -\log(S) (1 - \alpha)$ |
| **Mortality hazard rates; $h_1$ and $h_2$**<br>(Appendix S2 - this paper) | $P_1 = (1 - e^{-(\bar{h}_1 + \bar{h}_2)}) \frac{\bar{h}_1}{\bar{h}_1 + \bar{h}_2}$<br>$P_1 = (1 - e^{-(\bar{h}_1 + \bar{h}_2)}) \frac{\bar{h}_2}{\bar{h}_1 + \bar{h}_2}$<br>$S = e^{-(\bar{h}_1 + \bar{h}_2)}$ | $S = e^{-(\bar{h}_1 + \bar{h}_2)}$<br>$\alpha = \frac{\bar{h}_1}{\bar{h}_1 + \bar{h}_2}$ | $\bar{h}_1 = \text{'Time-averaged mortality hazard rate of cause 1'}$<br>$\bar{h}_2 = \text{'Time-averaged mortality hazard rate of cause 2'}$ |

<sup>1</sup> Parameters $P_1$ , $P_2$ and $S$ constrained to sum to 1 by e.g. the 'mlogit' link-function ( $P_k = \frac{\exp(\eta_k)}{1 + \sum_{l=1}^K \exp(\eta_l)}$ and $S = 1 - \sum_{k=1}^K P_k$ ).
<sup>2</sup> In the three references $\alpha$ was interpreted as 'Proportion died of cause 1' which does not assume that hazard rates remain proportional within intervals.

Since a cause-specific mortality probability *P*_*k*_ does not only depend on the mortality hazard rate for cause *k*, but also on the total mortality hazard rate (Appendix S2), it is often biologically more relevant and straightforward to model and interpret the cause-specific mortality hazard rates (*h*_*k*_’s). Nevertheless, statistical modelling of capture recapture data incorporating multiple (usually two) sources of mortality, has so far, to our knowledge, only been based on parameterizations for transition probabilities that do not include hazard rates explicitly (first two parameterizations in Table 1). Commonly used software for fitting multi-state capture-recapture data such as MARK (White & Burnham 1999) and E-SURGE (Choquet, Rouan & Pradel 2009) use the multinomial logit-link function to constrain the multinomial probability vector to sum to 1, which does not enable modelling of cause-specific hazard rates (Appendix S2). In the supplementary material we provide an R function that can be used with the ‘marked’ package (Laake, Johnson & Conn 2013) to model cause-specific hazard rates in capture-recapture analyses and give an example using simulated data in Appendix S3. This function can also be used to model additive effects on the total hazard rate. Even though methods for estimating cause-specific mortality hazard rates from time-to-event data have been more available (Lunn & McNeil 1995), the majority of ecological studies based on such data from telemetry or loggers also seem to put undue emphasis on comparing mortality probabilities rather than hazard rates.

Many studies have estimated relationships between probabilities of dying from different causes across years or space in order to assess whether mortality from these causes are additive (not associated), compensatory (negatively associated) or depensatory (positively associated) (Bischof *et al.* 2009; Servanty *et al.* 2010; Péron 2013; Wolfe *et al.* 2015). One problem of using mortality probabilities in such studies is that they are, as explained above, intrinsically related; increasing the mortality hazard rate of one cause implies that the probability of dying from this mortality cause will increase, while the probability of dying from all other causes decreases (Appendix S2). As shown in Figure 3, this intrinsic negative association may dominate empirical correlations between cause-specific mortality probabilities across years (or groups of individuals) even when correlations between the cause-specific mortality hazard rates across years (or groups) are strongly positive. Realizing this problem, researchers have come up with solutions to reduce “bias” [^1^ Note that correlations between mortality probabilities only are “biased” in the sense they do not represent correlations between cause-specific mortality hazard rates. They are not biased when interpreted strictly as correlations between interval-specific mortality probabilities.] in correlations between cause-specific mortality probabilities (Schaub & Lebreton 2004; Servanty *et al.* 2010) or to interpret these correlations with respect to hypotheses for compensatory versus additive mortality patterns (Péron 2013). However, when modelling mortality hazard rates, correlations can be interpreted directly; when partitioning mortality into one specific cause (e.g. “harvesting/predation”) and all other causes, a negative correlation between the mortality hazard rates among years indicates compensation, a positive correlation indicates depensation, and a correlation close to zero may indicate additivity. Furthermore, a near constant sum of the hazard rates indicates perfect compensation, while a negative correlation between the sum of the hazard rates and the harvesting hazard rate indicates overcompensation. When assessing such correlations it is important to ensure that the process correlations are not confounded with sampling correlations, for example by using appropriate hierarchical models (Royle & Dorazio 2008).

**Figure 3.**
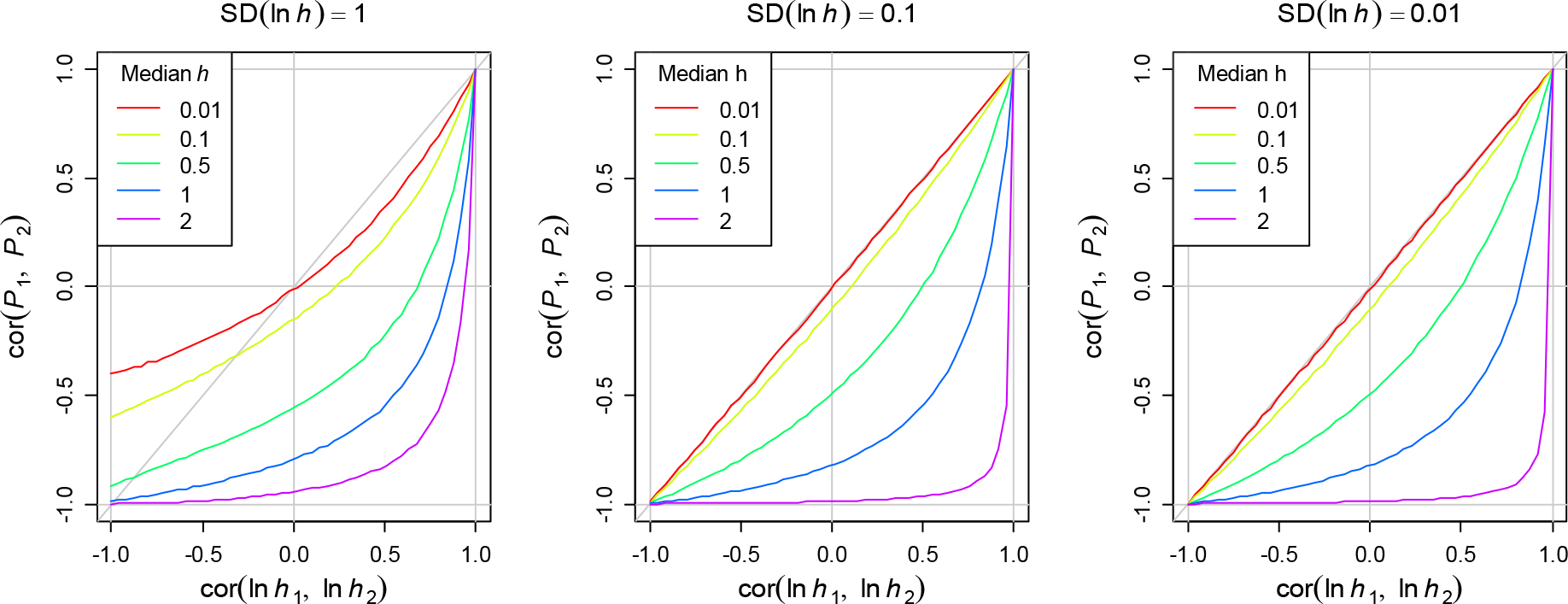
Correlations between probabilities of two mutually exclusive and collectively exhaustive causes of mortality (e.g. among years) as a function of the underlying correlation between the corresponding mortality hazard rates on a logarithmic scale (x-axes), assuming that the two hazard rates *h*_1_ and *h*_2_ have a bivariate log-normal distribution. Correlations were computed by first drawing 10^5^ values of ln(*h*_1_) and ln(*h*_2_) and then computing the Pearson correlation coefficients of *P*_1_ and *P*_2_ (Table 1). The standard deviation of ln(*h*_1_) and ln(*h*_2_) varies by panel as indicated above the plots. Different lines represent different median hazard rates according to colour legend.

Our recommendation to model and interpret mortality hazard rates instead of mortality probabilities is also highly relevant for studies of age related patterns in mortality. For instance, Koons *et al.* (2014) assessed patterns of senescence in a population of snow geese (*Chen caerulescens*) and a population of roe deer (*Capreolus capreolus*) by estimating age-dependent cause-specific mortality probabilities pertaining to hunting/human-related causes as well as natural causes. For both species they concluded that the yearly probability of dying from hunting/human-related causes was nearly independent of age, while the probability of dying from natural causes increased sharply at higher ages. However, as explained above, for the probability of dying from human-related causes to remain constant while the probability of dying from other causes increases with age, the mortality hazard rate representing human related causes must also increase to compensate for the loss of opportunities (Figure 4; note the >3 times increase in the hazard rate for the human related causes in the right panel).

**Figure 4.**
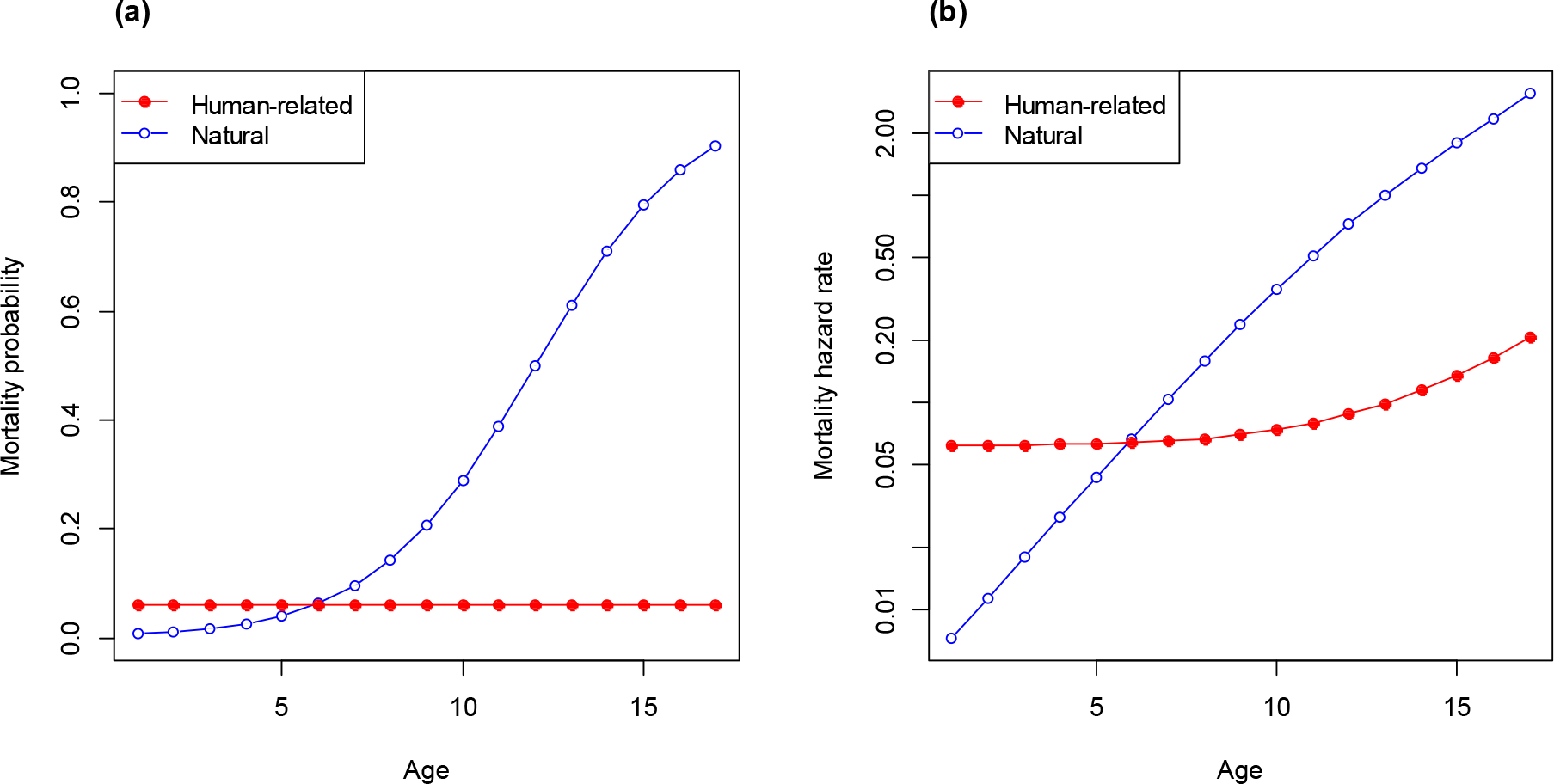
An example of two age-related competing causes of mortality; human-related and nonhuman-related (natural). Mortality probabilities are plotted in panel a, while corresponding time-averaged mortality hazard rates are shown in panel b. The fact that one of the two mortality probabilities is constant (red line in panel a) while the other increases with age, implies that both cause specific mortality hazard rates must increase with age (panel b) - see equations in Table 1. Mortality hazard rates (panel b) are plotted on a logarithmic scale for easier comparison of relative differences. In this example, the mortality hazard rate for human related causes (red line in panel b) increases more than 3 times from age 1 to age 17 even though mortality probability remains constant. Compare figure with Figure 4 in Koons *et al.* (2014).

### Mortality hazard rates and elasticity analyses

While statistical modelling of capture-recapture data dealt with so far aims to explain and predict variation in mortality rates and survival probabilities, we need a complete demographic model to assess the effects of these demographic parameters on population growth. Since there is often not sufficient data to fit a full demographic model, it is common to evaluate potential effects of changing demographic parameters and environmental variables using sensitivity analyses. Sensitivities are also essential in assessment of possible future environmental changes and in theoretical studies relating to population dynamics, management of natural populations, life-history theory and evolution (Benton & Grant 1999; van Tienderen 2000; Caswell 2001; Caswell 2007b).

When using discrete-time projection models it is common to evaluate the sensitivity of population growth rate (λ) to survival probabilities (*S*) in the model (as well as other parameters), either as the derivative *∂λ/∂λ* (sensitivity in the strict sense) or on relative scales, 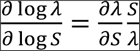 referred to as the elasticity of *λ* to survival probability. Given the fundamental nature of mortality hazard rates and the benefits of basing the statistical modelling of environmental effects on mortality hazard rates discussed in the above sections, it should be interesting to also evaluate sensitivities and elasticities of *λ* with respect to mortality hazard rates, to assess population level consequences of the effects. When log-linear effects for either the total or additive (cause-specific) components of the total hazard rate have been obtained, elasticities to hazard rates are particularly relevant as they are one component of the elasticities to the environmental variables. For example, if the effect on the hazard rate of an environmental variable *x* has been estimated in a log-linear model as 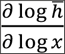 the contribution from this hazard rate to the total elasticity of *λ* to the environmental variable is 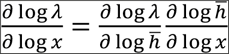 Even when such effects are not estimated, it may be reasonable to assume that e.g. a certain environmental change, management action or genetic change has a proportional effect on the mortality hazard rate. It can also be useful to compare elasticities to different ratio-scaled traits such as fertility, age at maturation and hazard rates (how would *λ* respond to a 1 percent decrease in fertility compared to a 1 percent increase in the mortality hazard rate?).

Recall that survival probabilities are defined by the step lengths of the discrete models, whereas mortality hazard rates represent the instantaneous risks of mortality and are invariant to any change in step length of the discrete model. If we for simplicity consider a case where the mortality hazard rate is constant over time in a birth-flow population, the elasticity of *λ* to a survival probability parameter defined for short intervals (e.g. monthly intervals) is naturally higher than the elasticity to a survival probability defined for long intervals (e.g., yearly intervals), whereas the elasticity to the mortality hazard rate is independent of the step length (Figure 5, Appendix S1).

**Figure 5.**
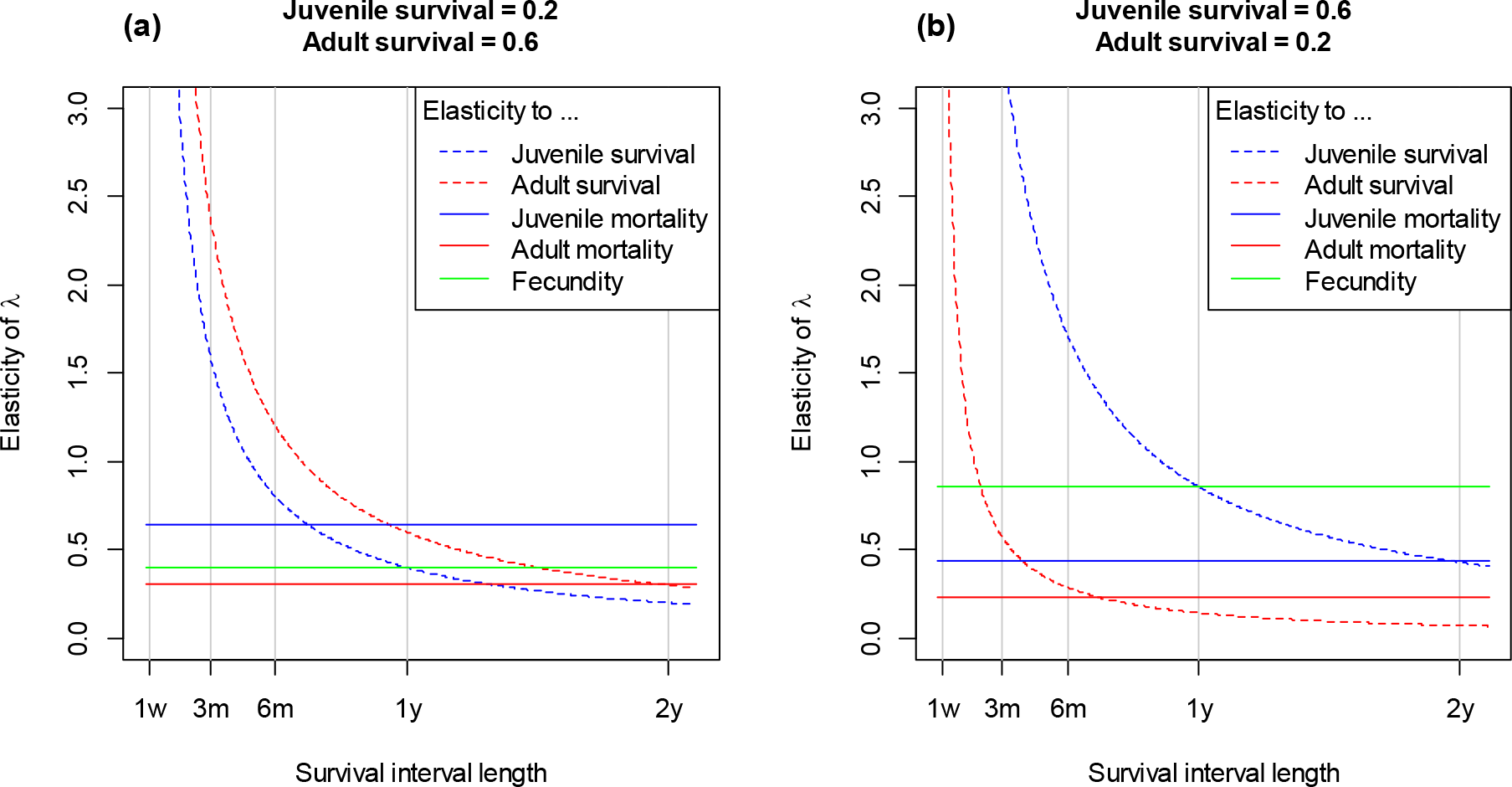
Elasticity of population growth rate (*λ*) to juvenile (blue lines) and adult (red lines) survival probabilities (dashed lines) and mortality hazard rate (solid lines), as well as to fecundity (green lines), when the model is parameterized with survival probabilities over different interval lengths (x-axis). The model assumes maturation at age 1 years, and fecundity is 2 offspring per year for all ages. Vertical lines indicate 1 week, 3 months, 6 months, 1 year and 2 years. In panel (a), yearly juvenile survival is 0.2, while adult survival is 0.6. In panel (b), yearly juvenile survival is 0.6, while adult survival is 0.2. For mortality hazard rates, the negative elasticity (i.e., the elasticity of lambda to a *decrease* in mortality hazard rate, or an increase in “current life-expectancy” (Appendix S1)) is plotted for easier comparison.

Since mortality hazard rates and survival probabilities are fundamentally different representations of the underlying mortality process, it should not be surprising that age specific patterns in the elasticities of *λ* to each parameter also differ (Figure 5). These patterns may even be diametrically different, highlighting the importance of clearly defining what representation of the mortality process is used in a given model to obtain valid biological inference. For example, in Figure 5a, where yearly juvenile survival probability is much lower than yearly adult survival probability (typical for iteroparous vertebrates), population growth rate has a higher elasticity to adult survival probability than to juvenile survival probability. However, elasticity with respect to mortality hazard rates shows the opposite pattern; elasticity to juvenile mortality hazard rate is higher than to adult mortality hazard rate. This result may seem contradictory if both these elasticities are loosely interpreted as representing the underlying mortality process in a similar way (which they do not). Figure 6a shows how elasticity ratios of *λ* with respect to survival probabilities relate to juvenile and adult survival and Figure 6b shows the equivalent elasticity ratios for mortality hazard rates. These very different patterns are due to the fact that a relative change in mortality hazard rate leads to a small relative change in survival probability when survival is high, but a much larger relative change when survival probability is low (Figure 7).

**Figure 6.**
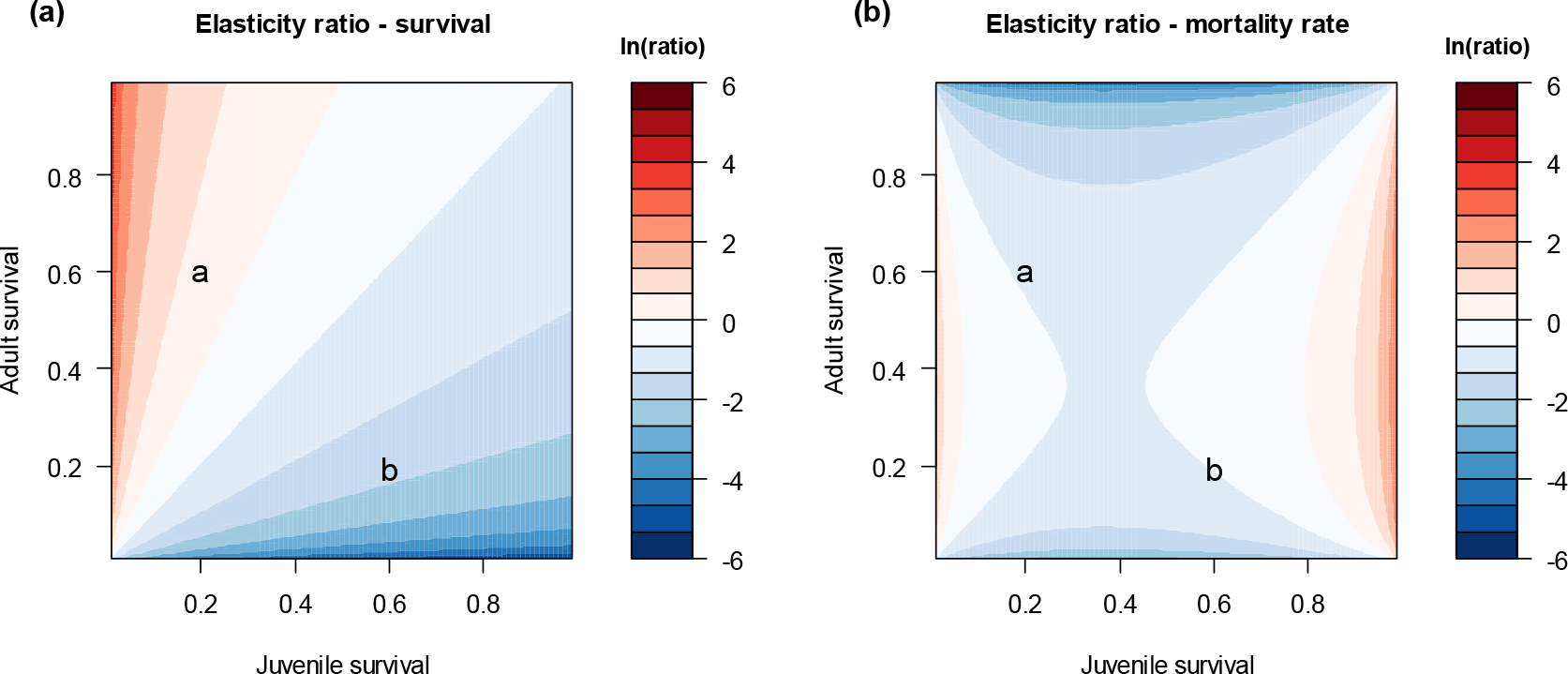
Elasticity of population growth rate (λ) to adult survival (panel a) and mortality hazard rate (panel *b*) relative to elasticity to respectively juvenile survival and juvenile mortality hazard rate. In both panels, the elasticity ratios (colour-key on a log_e_-scale) are shown for all possible combinations of juvenile (x-axis) and adult survival (y-axis). As in Figure 5, maturation occurs at age 1 and fecundity is 2 offspring per individuals for all ages. Plotted labels “a” and “b” correspond to the survival probabilities used in respectively panel (a) and panel (b) of Figure 5. It can be shown that the elasticity ratio with respect to mortality (panel b) is the hazard ratio 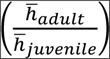 times the elasticity ratio with respect to survival (panel a).

**Figure 7.**
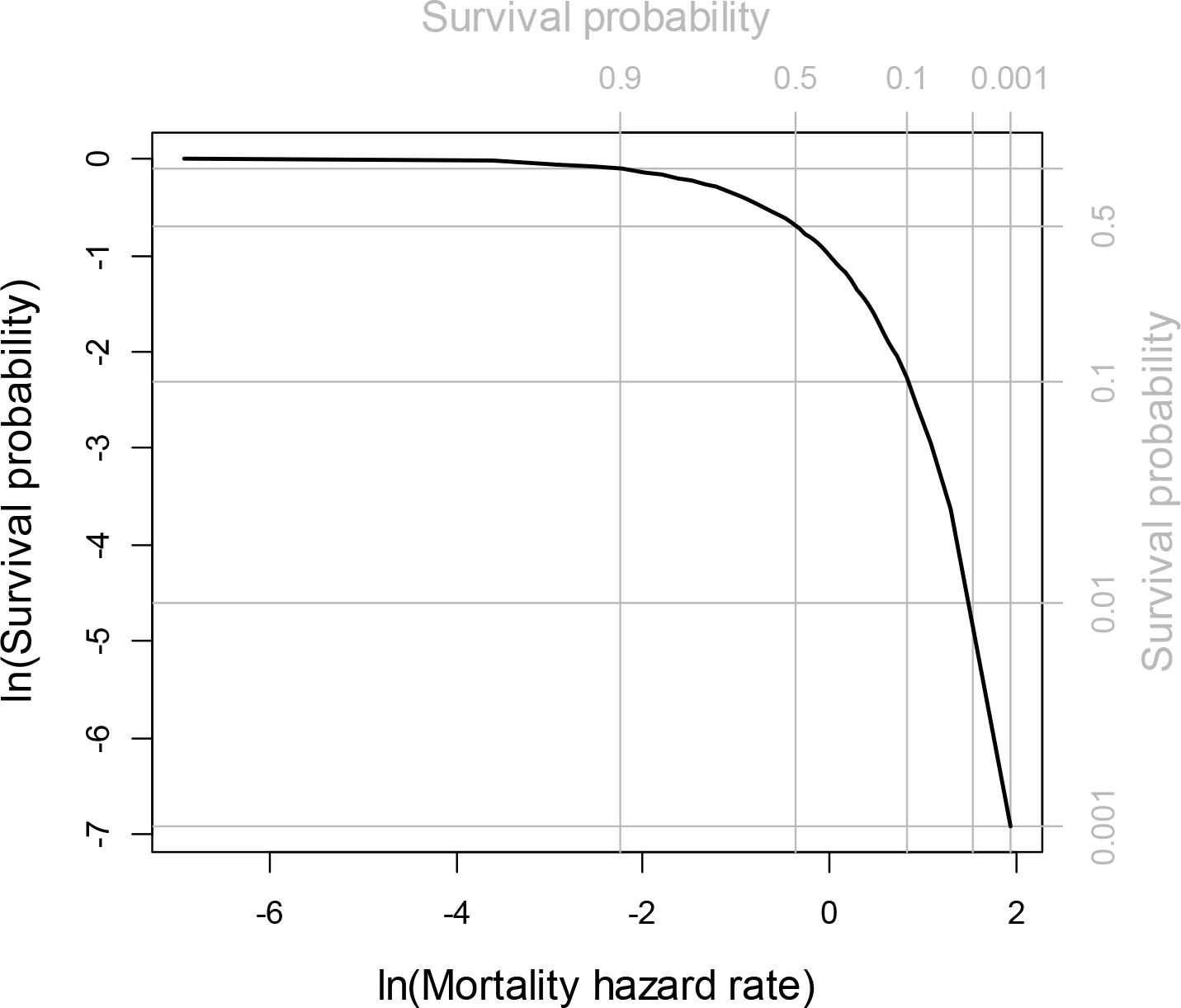
The natural logarithm of survival probability (y-axis) as a function of the natural logarithm of time-averaged mortality hazard rate (x-axis). The slope of the curve is the relative change in survival per relative change in mortality hazard rate. Corresponding survival probabilities are shown in the top and right axes and grey lines.

The fact that elasticities to different representations of the mortality process (as in Figure 6) may yield very different patterns has been noted in several comparative studies (see review in McDonald *et al.* (2017)). Motivated by the fact that variance in survival probability among years is limited by the mean (it cannot be as high when mean survival probability is low or high as when it is intermediate), McDonald *et al.* (2017) used the logit-transformation of survival probabilities as a variance-stabilizing transformation (Link & Doherty 2002) to compare patterns with respect to variability (on the transformed scale) and elasticity of demographic parameters in plant species. These patterns appeared very different than when using elasticities with respect to survival probabilities. Note that the loglog-transformation described before can also be seen as a variance-stabilizing transformation. Whereas the logit-transformation makes the variance independent of the mean when among-year variance in survival probability is proportional to 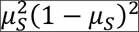 where *μ*_s_ is the mean survival among years, the loglog-transformation stabilize the variance when the variance is proportional to *μ*_*S*_^2^ log(*μ*_*s*_)^2^ (Appendix S1). Since among-year variability in survival probability is presumably highly system specific as it depends on complex trophic interactions and (non-linear) environmental effects relating to for example predators, pathogens, climate and intraspecific population density, it is difficult to argue that one variance-mean relationship (and transformation) is generally more applicable than others across systems. Note however that using the variance-stabilized sensitivity with respect to loglog-transformed survival probability is identical to the elasticity for mortality hazard rate (Appendix S1). As argued earlier, hazard-ratios (corresponding to the loglog-link) have a biologically more straightforward interpretation than odds-ratios (corresponding to the logit-link). Note also that there is no upper constraint for the variance of the mortality hazard rate among years.

Although we have here emphasized the utility of mortality hazard rates in elasticity analyses, we are not suggesting that it is wrong to use elasticities with respect to survival probabilities (or mortality probabilities, which would also yield different patterns; Link & Doherty 2002, Appendix S1). Appropriate measurements and scales always depend on application and assumptions. However, given the widely different patterns we have demonstrated here, we challenge practitioners to justify why they use for example elasticities to survival probabilities for specific interval lengths rather than (time-averaged) mortality hazard rates in their analyses. Finally, we note that there is no technical reason for preferring the use of elasticities with respect to probabilities as the corresponding elasticities with respect to mortality hazard rates can easily be computed with a simple application of the chain-rule of derivation (the R code give in Appendix S1 used to produce Figures 5 and 6 makes use of a function in the ‘popbio’ package (Stubben & Milligan 2007) and can be used for any matrix projection model).

### Conclusion and perspectives

Mortality is a process in continuous time, and mortality hazard rates are the most fundamental measurements describing this process. We have here highlighted the benefits of basing inference on hazard rates, also in discrete-time models. We recommend modelling log-linear (multiplicative) effects on time-averaged hazard rates by using a loglog-link function for survival probabilities. In models involving multiple causes of death, estimates of hazard rates can be obtained by using a hazard rate based multinomial link function, assuming that the cause-specific hazard rates remain proportional within intervals (Table 1). This allows direct inference regarding interactions between different sources of mortality. When the assumption about proportionality within intervals is not reasonable, the intervals may split up in such a way that e.g. hunting mortality can only occur during the hunting season.

The issues we have discussed here apply more generally to any process involving transitions that can take place instantaneously in continuous time, such as transitions between disease states (Conn, Cooch & Caley 2012) or reproductive states of multivoltine species (Ergon, Yoccoz & Nichols 2009). When more than one transition can take place at any point in time within the intervals, an increase in the hazard rate for one transition will reduce the probability that the other competing transitions will occur. As a general method, transition probability matrices may be computed from matrices of transition hazard rates by the use of matrix exponentials (Keyfitz & Caswell 2005 chap. 17.5; Miller & Andersen 2008; Conn, Cooch & Caley 2012). These models may in principle include intermediate and reversible transitions, but identifiability of statistical models can be an issue. It is also often desirable to focus the modelling on the latent probability density function for time/age of state-transition rather than the corresponding hazard rate (Ergon, Yoccoz & Nichols 2009; de Valpine *et al.* 2014; Appendix S1 1.4.1.4 and 1.4.1.5).

Methods focusing on the modelling of hazard rates are well established and much used in other fields, but seem to a large extent to be lacking from ecologists’ toolbox. This may in part be due to lack of attention to meaningful inference (Houle *et al.* 2011) as well as tradition and the fact that hazard rate based models (other than loglog-link models of survival probabilities) have not yet been implemented in software commonly used by ecologists. We conclude that many areas of ecological research would benefit from an increased awareness of the utility of transition hazard rates in modelling and inference.

## Acknowledgements

We greatly appreciate comments by Hal Caswell, Guillaume Péron, Nigel G. Yoccoz and one anonymous reviewer on earlier versions of the manuscript.

## Authors’ contributions

TE conceived the main ideas and wrote the first version of the manuscript. All authors contributed with discussions, literature review, commenting and editing of subsequent versions.

## Data accessibility

This article does not use collected data. R code for producing all figures and other supporting material is provided in the online version of the article.

## Box 1 Mortality hazard rates and survival probabilities

**Figure B1.**
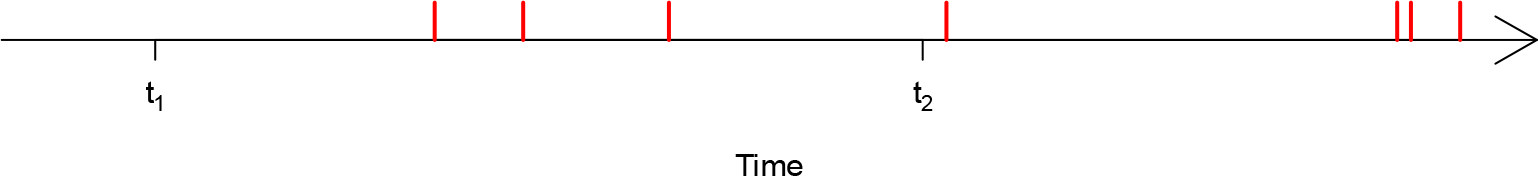
Realized times of recurrent events (or “deadly events experienced by an immortal individual”)

A mortality hazard rate is the underlying intensity of deadly events that an individual is exposed to. To better understand the relationship between survival probabilities and hazard rates, first consider a point process of non-lethal events (e.g., ingestion of food items). An example of realized times of such events for an individual is shown in Figure B1. Denoting the intensity of this point process as a function of time as *λ*(*t*), the expected number of events occurring during an interval from time *t*_1_ to time *t*_2_ is the integrated intensity over this interval, 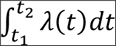. When events occur independently of each other, the number of events *X* occurring in the interval is a random Poisson variable, *X*~Poisson 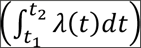. If these events are lethal, *X* must be zero for the individual to survive the interval, which according to the Poisson distribution occurs with probability 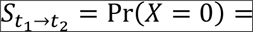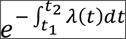. When studying lethal events, the event intensity is called a ‘hazard rate’ and is often denoted as *h*(*t*) (= λ(*t*) above). A calculus-based derivation of the relationship between hazard rates and survival is provided in Appendix S1.

Mortality hazard rates may thus be interpreted as the “expected number of deadly events occurring per time-unit for an immortal individual”. Since hazard rates are **ratio scaled** intensities, they may be expressed with an arbitrary time-unit and values are proportional to the length of the time-unit used (e.g, a hazard rate of one event per month is equivalent to 12 events per year). The ratio of two **hazard rates**, a hazard ratio, is thus invariant to the time-unit used to express the rates and has a clear interpretation; if the mortality hazard rate for juveniles is twice the mortality hazard rate of adults, we know that juveniles are exposed to twice as many expected deadly events per time-unit than adults. Such hazard ratios may be estimated by modelling survival probability through a **loglog-link**; *S* = exp(—exp(*η*)), where *η* is the linear predictor and exp(*η*) is the time-averaged mortality hazard rate over the interval, expressed with the interval length as a time-unit. If survival probability of one group of individuals is represented by *η* = *β*_0_ + *β*_2_log(*x*) and another by *η*_2_ = *β* + β_1_ + *β*_2_log(*x*), where *x* is a continuous covariate, the hazard ratio representing their **relative difference** in mortality, irrespective of the covariate, becomes 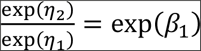. Further, when the covariate is multiplied by a factor *C*, the mortality hazard rate will be multiplied by a factor 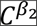 (when *β*_2_ = 1 the hazard rate is proportional to the covariate). Hence, when using a loglog-link on survival probabilities we model relative differences in hazard rates (i.e., hazard ratios). [Note that some computer programs like MARK (White & Burnham 1999) define the loglog-link function as *S* = exp(—exp(—*η*)), in which case the above hazard ratio becomes exp(—*β*_1_).]

### Glossary

**Mortality hazard rate:** The latent intensity of deadly events that an individual is exposed to (Box 1). Synonymous to ‘force of mortality’ and ‘failure rate’. Often referred to as just ‘mortality rate’.

**Survival probability**: The probability that an individual survives an interval of a given length given that it is alive at the beginning of the interval. A unit-free measurement on an absolute scale. Often misleadingly referred to as ‘survival rate’.

**Mortality probability**: The probability that an individual dies during a given time interval, equal to one minus the survival probability for the same interval. Often misleadingly referred to as ‘mortality rate’.

**Odds**: When applied to survival, **survival probability** divided by **mortality probability**.

**Odds-ratio**: One odds divided by another.

**Hazard ratio**: One hazard rate divided by another.

**Logit-link**: A link-function on the form log(‘odds’) = ‘linear predictor’ =*β*_0_ +**β**_1_*x*_1_ + *β*_2_*x*_2_ + … = ***βx***. Generally used for probabilities that are not survival probabilities.

**Loglog-link**: When applied to survival probability, a link-function on the form log(’time-averaged mortality hazard rate’) = ‘linear predictor’ (see Box 1).

**Permissible transformation**: Transformations that preserve the relevant relationships among measurements (Houle *et al.* 2011)

**Absolute-scaled variable**: A scale type with no **permissible transformations** (e.g. probabilities).

**Ratio-scaled variable**: A positive variable with a true (nature-given) zero (zero refers to “non-existence” or “total absence”). Examples include age, mass and mortality hazard rates. The only **permissible transformations** are multiplications and divisions by strictly positive numbers.

*Absolute difference/effect*: A difference or effect quantified as one quantity minus another. E.g., the absolute difference between values 2 and 1 is 1.

**Relative difference/effect**: A difference or effect quantified as one quantity divided by another. E.g., the relative difference between values 2 and 1 is 2. The **absolute difference** on a logarithmic scale equals the logarithm of the relative difference since log(a) − log (*b*) = log(*a/b*).

**Linear contrast**: The **absolute difference** between two predicted values at the scale of the link-function.

**Log-linear models**: Models on the form log(‘prediction’) = *β*_0_ +*β*_1_*x*_1_ + *β*_2_*x*_2_ + …. A common choice for **ratio-scaled variables**.

## Appendix S1: Details on sections and figures

The headings below are the same as in the article. Each section contains explanations and R-code for the figures referred to under each heading.

## 1.1 Introduction (Box 1)

## 1.1.1 The relationship between mortality hazard rates and survival probabilities

In Box 1 we explained the relationship between individual mortality hazard rates and survival probabilities (eq. (1) in the main text) by considering an individual point process for deadly events. We chose this approach to give an intuitive understanding of the hazard rate as an “intensity of deadly events” that individuals are exposed to and to set the stage for explaining the issue of competing risks in an intuitive manner in Appendix S2. The following derivation follows the derivation in Quinn & Deriso (1999, p. 10), referring to Ricker (1975), only that we here treat the mortality hazard rate as an individual trait and not a population level characteristic.

Consider a hypothetical population of *N*(*t*_1_) identical individuals having identical expectations for their future environment at time *t*_1_. If there is no reproduction, the instantaneous rate of change in number of individuals, *N*(*t*), will be

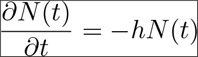

where *h* is the mortality hazard rate in units of *t*^−1^, here considered to be constant. Letting Δ represent a certain increment in time and solving for *N*(*t*_1_ + Δ) as a function of *N*(*t*_1_) gives

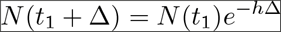

To convert this to a survival probability over the interval, we divide by *N*(*t*_1_),

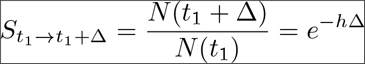

If the mortality hazard rate is considered as a continuous function of time, *h*(*t*), we can calculate the survival probability as a product of many infinitesimally small sections of the interval from time *t*_1_ to *t*_2_ to obtain eq. (1) in the main text,

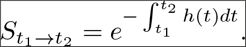

For further relationships and derivations, see Collet (2014, chap. 1.3) and section 1.4.1.5 below.

As an analogy, the safety of a car or driver may be described as the expected number of collisions per kilometre (i.e., a hazard rate). This hazard rate may be expressed with any distance unit. One can also express the safety of the car or driver as a probability of avoiding collision (i.e., a survival probability) when driving a given distance. A probability has no unit, so the distance for which the probability is defined is essential for its meaning. If the probability is defined at a very short distance the probability of avoiding collision is almost one, and if the distance is long enough, the “survival” probability is almost zero. Even if substantial road (environmental) improvements were made, these two probabilities would still be approximately one (infinite odds) and zero (zero odds), irrespective of the driver (individual differences). When modelling environmental (road) effects, a natural starting point is to assume that environmental differences (road improvements) have a proportional effect (i.e., a certain percentage change in the hazard rate). This leads to a log-linear model for the hazard rate (see section on “Log-linear models of mortality hazard rates vs. logit-linear models of survival probabilities”). Alternatively, if the total hazard rate is composed of several cause-specific hazards, and some of these causes are eliminated, a model with additive effect of the total hazard rate would be appropriate (see section on “Modelling of multiple sources of mortality and competing risks”). A proportional (log-linear) or additive model for survival probabilities or odds defined for a *predefined* driving distance (or time) requires a justification for the choice of driving distance used in the model, which seems more arbitrary.

## 1.2 Log-linear models of mortality hazard rates vs. logit-linear models of survival probabilities

## 1.2.1 Relationships

Survival probabilities, *S*, in any discrete-time model can be replaced by *S* = exp(−*h*̄), where *h̄* is the time-averaged mortality hazard rate in the given time interval(s). The unit of measurement for *h̄* is here the inverse of the length of the interval (see Box 1 in the main text). That is, if *S* is the probability of surviving one year, the unit of *h̄* is year^−1^, whereas if *S* is the probability of surviving one month, the unit of *h̄* is month^−1^.

Since *h̄* is a ratio-scaled intensity, it is natural to model multiplicative (proportional) effects on *h̄* through a log-link,

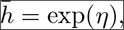

where *η* is a linear predictor of explanatory variables,

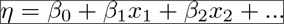

This corresponds to using a loglog link-function on survival probabilities,

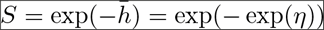

The exponent of contrasts in the linear predictor (e.g. representing different age groups or individuals with different body mass or sex) then becomes hazard ratios [^i^ Note that some computer programs for analyses of capture-recapture data, such as MARK, define the inverse loglog-link as *S* = exp(−exp(−*η*)), in which case the hazard ratio 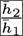 becomes exp(*η*_1_ − *η*_2_).],

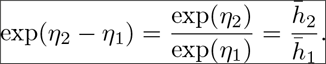

One could reparameterize the discrete-time model with e.g. monthly survival probabilities (*S*_*m*_) instead of yearly survival probabilities (*S*_*y*_). If one assumes that all months of the year have the same survival probability, this amounts to just replacing all *S*_*y*_ in the model with (*S*_*m*_)^12^. [^ii^ Note that this is a reparameterization since *S*_*y*_ and *S*_*m*_ have different meanings - it is not about “selecting a time-scale for the survival probabilities”. Survival probabilities are unit-free absolute scale measurements that cannot be “re-scaled” in any way. Such reparameterizations of the general (full, unconstrained) model may be specified in e.g. MARK, and is commonly done when time between capture occasions vary over the study period.] When using a loglog-link on survival probabilities, the parameters in the model is invariant to such replacements because the time-averaged hazard rate remains unchanged. If this does not appear as obvious, one may see this better by first calculating the time-averaged hazard rate as *h̄*_*y*_ = − log(*S*_*y*_). The unit for *h̄*_*y*_ is here year^−1^. Similarly, the time-averaged hazard rate can also be calculated as *h̄*_*m*_ = − log(*S*_*m*_), where *h̄*_*m*_ is expressed in units of month^−1^. Since *S*_*y*_ = (*S*_*m*_)^12^, we see that *h̄*_*y*_ = − log((*S*_*m*_)^12^) = —12log(*S*_*m*_) = 12*h̄*_*m*_, and since an intensity of one event per month is the same as an intensity of 12 events per year, the two hazard rates *h̄*_*y*_ and *h̄*_*m*_ are identical (they just have different units of expression). Note also that the ratio of two hazard rates (i.e., hazard ratios) are invariant to which unit of expression is used for the hazard rates. Hence, when modelling survival probabilities through a loglog-link, all parameters in the linear predictor except the intercept are invariant to reparameterizations on the form *S*_*y*_ = (*S*_*m*_)^12^. This can be seen from expanding *h̄*_*y*_ = 12*h̄*_*m*_,

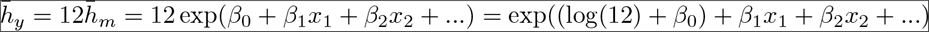

I.e., the effect sizes in the log-hazard model is “parameterization invariant” (Pace & Salavan 1997).

It is (unfortunately) common practice to model effects of covariates on survival probabilities through the use of a logit link-function. That is, survival probability is modelled as

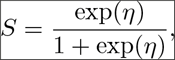

where *η* is the linear predictor. The exponent of the linear predictor then becomes the survival-odds,

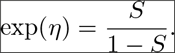

Just like survival probabilities, but unlike hazard rates, survival-odds are unit-free absolute scale measurements that cannot be transformed in any way without losing their original meaning.

If we fit an effect of a time-invariant covariate (e.g., an individual covariate) on monthly survival using a logit-link, the relationship between the covariate and yearly survival will be

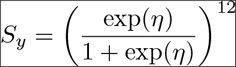

which is not easily simplified. Hence, replacing *S*_*y*_ with (*S*_*m*_)^12^ changes the functional form of the model when survival probabilities are modelled by a logit-link involving continuous covariates. In contrast, when using a loglog-link, we obtain

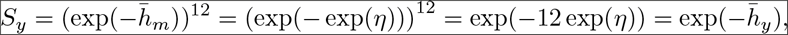

and, as also explained above, the functional relationships between the covariates and survival remains unchanged (only the intercept is changed to reflect the change in unit of expression for the time-averaged hazard rates).

In conclusion, when using the model *S* = (logit^−1^(*β*_0_ + *β*_i_*x*_1_))^*n*^ where *n* is the length of the survival interval in the chosen time-units, *n* can be interpreted as a parameter that is fixed by the modeller (e.g., by setting the time-interval length in MARK). In contrast, the model *S* = (loglog^−1^*β*_0_ + *β*_1_*x*_1_))^*n*^ is equivalent to 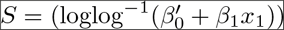 where 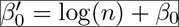. Hence, when using a loglog-link, changing the time-scale just amounts to changing the definition of the intercept parameter but the model remains the same (i.e., it is a reparameterization).

## 1.2.2 Figure 1

Figure 1 in the main text illustrates the effect of replacing *S*_*y*_ with (*S*_*n*_)^*n*^ where 1/*n* is the length of the time-intervals used in the discrete model. To facilitate comparison of functional shapes, the intercepts and slopes of logit survival for all curves are defined such that yearly survival probability equals 0.05 when the covariate value is −2 and 0.95 when the covariate value is 2. If the models were fitted to data, the functional forms would be the same as in the figure, and predicted values would be in the same range, but the curves would not necessarily cross at the same place.

## 1.2.2.1 Code - Figure 1

~~~
*# Defining functions*
logit = function(S) log(S/(1-S))
invlogit = function(eta) exp(eta)/(1+exp(eta))
loglog = function(S) log(−log(S))
invloglog = function(eta) exp(−exp(eta))
rescaled.invlogit = function(x, il, s1=0.05, s2 = 0.95){ *# il = 'interval length'*
  a = mean(c(logit(s2^il), logit(s1^il))) *# intercept*
  b = (logit(s2^il)-logit(s1^il))/4 *# slope*
  Sy = invlogit(a + b*x)^(1/il) *# yearly survival*
  return(Sy)
}
rescaled.invloglog = function(x, il, s1=0.05, s2 = 0.95){ *# il = 'interval length'*
  a = mean(c(loglog(s2^il), loglog(s1^il))) *# intercept*
  b = (loglog(s2^il)-loglog(s1^il))/4 *# slope*
  Sy = invloglog(a + b*x)^(1/il) *# yearly survival*
  return(Sy)
}

*# Values for plotting*
x = seq(−3,3,length.out=60)
Syy.logit = rescaled.invlogit(x, 1) *# Using logit-link applied to yearly survival*
Sym.logit = rescaled.invlogit(x, 1/12) *# Using logit-link applied to monthly survival*
Sy12.logit = rescaled.invlogit(x, 12) *# Using logit-link applied to 12-year survival*
Syy.loglog = rescaled.invloglog(x, 1) *# Using loglog-link applied to yearly survival*
Sym.loglog = rescaled.invloglog(x, 1/12) *# Using loglog-link applied to monthly survival*

*# All Syy.loglog == Sym.loglog (but round-off errors occur, hence using round())*
all(round(Syy.loglog, 12) == round(Sym.loglog, 12))

## [1] TRUE
*# Making plot*
plot(x, Syy.logit, type="l", col="blue",
     xlab="Time-invariant covariate", ylab="Yearly survival")
lines(x, Sym.logit, col="red")
lines(x, Sy12.logit, col="green")
lines(x, Syy.loglog, lty=2)
abline(v=-2, lty=3)
abline(v=2, lty=3)
legend("topleft",
  legend = c(
    expression(logit(italic(S)[monthly])),
    expression(logit(italic(S)[yearly])),
    expression(logit(italic(S)[12-yearly])),
    expression(loglog(italic(S^{n})))
 ),
  lty=c(1,1,1,2), col=c("red", "blue", "green", "black"), bg="white"
)
~~~

## 1.2.3 Figure 2

Odds-ratios are hard to interpret, not only because their meaning depend on the chosen interval length of the discrete-time model, but also because, for a given hazard ratio, they depend on the survival probability of the reference group. This is illustrated in Figure 2 in the main text.

## 1.2.3.1 Code - Figure 2

~~~
*# Function for computing odds-ratio for a given hazard ratio (hr) and survival*
*# probability of reference group (s1)*
OR = function(s1, hr){
  s2 = exp(−(−log(s1)*hr))
  (s2/(1-s2))/(s1/(1-s1))
}
ORv = Vectorize(OR)

*# Constants*
scale = seq(0, 2.1, length.out=60) *# Survival interval length in years (x-axis)*
s = c(0.01, 0.1, 0.25, 0.5, 0.75, 0.99) *# Survival probabilities of reference*
HR = 0.5 *# Hazard ratio to be used in plot*
*# Draw the plot*
Col = rainbow(length(s))
Col[2] = "#FF8000FF" *# Changing the light yellow to orange*
plot(1,1, type="n", ylim=c(0,6), xlim=range(scale), axes=F,
     xlab="Survival interval length", ylab="Odds-ratio")
box()
axis(2)
At = c(7/365, 0.5, 1, 2)
Lab = c("1w", "6m", "1y", "2y")
axis(1, at = At, labels=Lab)
abline(v = At, col="grey")
abline(h=1, lty=2)
lines(range(scale), c(HR,HR), lwd=2) *# Line for the hazard ratio*
for(i in 1:length(s)){
  lines(scale, ORv(s[i]^scale, HR), col=Col[i], lwd=1.5)
}
legend("topleft", legend=s, col=Col, lty=1, bg="white", lwd=1.5,
      title=" Yearly survival of reference ", ncol=2)

*# Adding points corresponding to the examples in the text*
points(c(1,2), ORv(c(0.25, 0.25^2), HR), cex=1.7) *# Original environment; odds-ratio*
points(c(1,2), c(HR,HR), cex=1.7) *# Original environment; hazard-ratio*
points(c(1,2), ORv(c(0.5, 0.5^2), HR), pch=4, cex=1.7)*# Improved environment; odds-ratio*
points(c(1,2), c(HR,HR), pch=4, cex=1.7) *# Improved environment; hazard ratio*
~~~

From Fig. 2, it may be noted that odds-ratios approach the inverse of the hazard ratio (1/0.5 = 2) when survival probabilities are high; i.e., when the survival probability of the reference is high (magenta line) or when interval lengths are short (to the left in the plot).

## 1.3 Modelling of multiple sources of mortality and competing risks

Mortality hazard rates are more elucidating with respect to the underlying mechanisms than mortality probabilities when studying associations between two or more sources of mortality in a population because mortality probabilities are intrinsically dependent; increasing the hazard rate for one cause of mortality will automatically reduce the probabilities of dying from all other causes (Appendix S2). Figure 3 in the main text shows how this negative intrinsic dependency can dominate empirical correlations between cause-specific mortality probabilities across years even when correlations between the cause-specific mortality hazard rates are strongly positive.

Figure 4 illustrates the relationships between cause-specific mortality hazard rates and survival probabilities in a age-dependent model. We note here that, in empirical studies of senescence, one should also consider potential biases arising from un-modelled individual heterogeneity (i.e., “frailty” [^iii^ Unobserved individual heterogeneity in mortality hazard rates. In the presence of frailty, the population will be increasingly dominated by individuals with low mortality hazard rates at higher age-classes. Hence, age-related patterns in population level mortality rates are not representative of individual level patterns.]) (Vaupel, Manten & Stallard 1979; Cam et al. 2002; Collett 2014), temporary emigration (Langtimm 2009) and age-dependent dispersal (Ergon & Gardner 2014).

## 1.3.1 Figure 3

Figure 3 is produced by simulation. First, 10^5^ values of log hazard rates, log(*h*_1_) and log(*h*_2_), were drawn from a bi-variate normal distribution with means given by the the colour legend (median(*h*) = exp(mean(log(*h*)))), standard deviations given above each panel, and correlations given by the x-axis. Cause-specific mortality probabilities where then computed from each of the 10^5^ samples, and the Pearson correlation coefficients between them where then computed and plotted on the y-axis.

## 1.3.1.1 Code - Figure 3

~~~
library(MASS) *# for ‘mvrnorm’*

*# Functions*
sim.cor = function(sigma, mu){
    Rho = seq(−1,1,length.out=50)
    cor.h = cor.P = median.S = rep(NA, length(Rho))
    for(i in 1:length(Rho)){
        rho = Rho[i] *# correlation between log hazards*
        S = matrix(c(
           sigma[1]^2, rho*sigma[1]*sigma[2],
           rho*sigma[1]*sigma[2], sigma[2]^2), 2,2, byrow=T)
        log.h = mvrnorm(100000, c(mu,mu), S)
        h = exp(log.h)
        *# Cause-specific mortality probabilities from Table 1 in main text:*
        P1 = (1-exp(−(h[,1]+h[,2])))*h[,1]/(h[,1]+h[,2])
        P2 = (1-exp(−(h[,1]+h[,2])))*h[,2]/(h[,1]+h[,2])
        cor.P[i] = cor(P1,P2)
   }
   return(list(Rho=Rho, cor.P=cor.P))
}

plot.panel = function(sigma){
  H = c(0.01, 0.1, 0.5, 1, 2)
  Mu = log(H)
  Col = rainbow(length(Mu))
  plot(0,0, type=“n”, xlab=expression(cor(log~italic(h[1]),~log~italic(h[2]))),
       ylab=expression(cor(italic(P[1]),~italic(P[2]))))
  abline(0,1, col=“grey”)
  abline(v=c(−1,0,1), col=“grey”)
  abline(h=c(−1,0,1), col=“grey”)
  for(i in 1:length(Mu)){
    X = sim.cor(rep(sigma,2), mu=Mu[i])
    lines(X$Rho, X$cor.P, col=Col[i])
  }
  legend(“topleft”, lty=1, col=Col, legend=H, title=expression(Median~italic(h)),
        bg=“white”)
}

*# Make each panel*
par(mfrow=c(1,3))
plot.panel(1)
title(expression(SD(log~italic(h)) == 1))
plot.panel(.1)
title(expression(SD(log~italic(h)) == 0.1))
plot.panel(.01)
title(expression(SD(log~italic(h)) == 0.01))
−1.0
~~~

## 1.3.2 Figure 4

Figure 4 illustrates the fact that, if one of two cause-specific mortality *probabilities* is constant (red line in panel a) while the other increases with age, both cause-specific mortality *hazard rates* must increase with age (panel b).

## 1.3.2.1 Code - Figure 4

~~~
*# Parameters chosen to resemble Fig. 4 in Koons et al. 2014*
x = 1:17 *# Age (x-axis values)*
c = 12 *# Age at 50% survival*
b = .45 *# Slope*

*# Cause-specific mortality probabilities*
P1 = rep(0.06, length(x)) *# Human related*
P2 = 1/(1+exp(−(b*(x-c)))) *# Natural*

*# Corresponding time-averaged mortality hazard rates from Table 1 in main text*
m1 = −log(1-P1-P2)*P1/(P1+P2)
m2 = −log(1-P1-P2)*P2/(P1+P2)

*# Plotting the two panels:*
par(mfrow=c(1,2))

plot(mean(x),mean(P2), xlim=range(x), ylim=c(0,1), type=“n”,
     ylab=“Mortality probability”, xlab=“Age”)
lines(x,P2, col=“blue”)
points(x,P2, col=“blue”, pch=21, bg='white')
lines(x,P1, col=“red”)
points(x,P1, col=“red”, pch=19)
legend(“topleft”, legend=c(“Human-related”, “Natural”), col=c(“red”,“blue”),
       pch=c(19,21), pt.bg = 'white', lty=1)
title(“(a)”, adj=0)

plot(mean(x),mean(m2), xlim=range(x), ylim=range(c(m1,m2)), type=“n”, log=“y”,
     ylab=“Mortality hazard rate”, xlab=“Age”)
lines(x,m2, col=“blue”)
points(x,m2, col=“blue”, pch=21, bg='white')
lines(x,m1, col=“red”)
points(x,m1, col=“red”, pch=19)
legend(“topleft”, legend=c(“Human-related”, “Natural”), col=c(“red”,“blue”),
       pch=c(19,21), pt.bg='white', lty=1)
title(“(b)”, adj=0)
~~~

## 1.4 Mortality hazard rates and elasticity analyses

## 1.4.1 Relationships

## 1.4.1.1 Elasticity to *h* and *S*

To derive the relationship between elasticity of population growth rate λ to survival probability *S* over an interval, *ϵ*_*S*_, and elasticity to time-averaged mortality hazard rate *h̄* during the same interval, *ϵ*_*h̄*_, note first the relationship *h̄* = − log(*S*) (eq. (2) in the main text) where the time-unit for *h̄* is the inverse of the length of the interval. By using the chain rule of derivation, we get

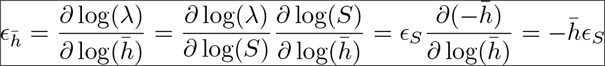

## 1.4.1.2 Dependencies on interval length

If the above survival probability applies to one year, *S*_*y*_, one may choose to reparameterize the model with monthly survival probabilities, *S*_*m*_, assuming that all months of the year have the same survival probability such that 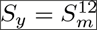 and log(*S*_*y*_) = 12log(*S*_*m*_). The relationship between the elasticities of these two survival probabilities is

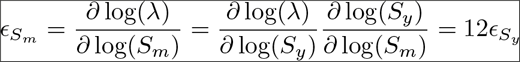

Elasticities to instantaneous mortality hazard rates, or their time-averaged values (*h̄* = − log(*S*)), are invariant to any such reparameterizations with respect to the choice of interval length of the discrete model. If this is not intuitive, consider the relationship 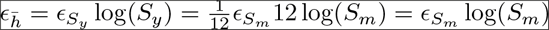.

## 1.4.1.3 Elasticities to additive (cause-specific) components of the total hazard rate

As explained in Appendix S2, when there are multiple causes of death, the total mortality hazard rate is the sum of the cause-specific hazard rates *h*_*k*_

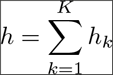

Applying the chain rule to find the elasticity to a cause-specific hazard rate, we get

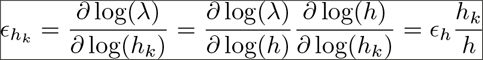

Hence, the elasticity to the total mortality hazard rate equals the sum of the elasticities to the cause-specific hazard rates,

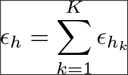

## 1.4.1.4 Elasticity to mortality probability

As pointed out by Link & Doherty (2002), the relationship between elasticity to mortality probability, *ϵ*_*P*_, and elasticity to survival probability, *ϵ*_*S*_, is

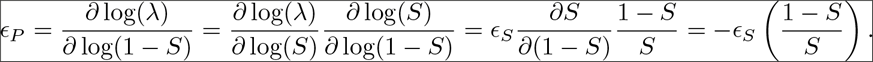

That is, a given relative decrease in morality probability only has the same impact on population growth rate (or fitness) as the same relative increase in survival probability when *S* = 0.5. Moving away from *S* = 0.5 in either direction, these two elasticities will be increasingly different and hence represent very different degrees of change in the underlying mortality process when *S* is not close to 0.5. Elasticity to mortality hazard rate, on the other hand, has a clear and meaningful interpretation; it is the relative change in population growth rate caused by a relative increase in the intensity at which individuals die (Box 1 in the main text). Further, as explained below, −*ϵ*_*h̄*_ is the elasticity to “current life-expectancy”.

## 1.4.1.5 Elasticity to “current life-expectancy”

The relationship between the mortality hazard rate *h*(*t*) and the probability density for death at time *t*, *f*(*t*), is given as

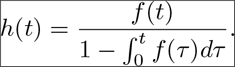

(Collet 2014, chap. 1.3). The probability density function *f*(*t*) is here conditional on being alive at *t* = 0 (defined as time of birth or the beginning of a given interval) and integrates to 1 over infinite time. The hazard rate function must always be greater than or equal to zero and must integrate to infinity over infinite time. It is also useful to note that the denominator in the above expression is the probability of being alive at time *t*, 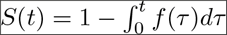. Further, the numerator of the expression is the negative derivative of this survival function. Hence, the hazard rate function can also be expressed as

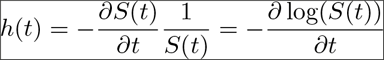

(Collet 2014, chap. 1.3).

Hence, for any given probability distribution for time of death *f*(*t*), there is a corresponding hazard rate function *h*(*t*), and vice versa. The probability distribution for time of death corresponding to a constant hazard rate *h*(*t*) = *h* is the exponential distribution with mean 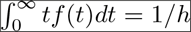. Hence, if an individual (hypothetically) maintains its current mortality hazard rate throughout life, its expected time until death (i.e., its “current life-expectancy”) is 1/*h*. The elasticity of λ to current life-expectancy is then

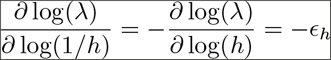

## 1.4.1.6 Perspectives on hazard rates and models of stage-duration distributions

The notion of elasticity to life-expectancy introduced in section 1.4.1.5 above is similar to the notion of elasticity to stage-duration in the absence of mortality (de Valpine et al. 2014). Structured population models based on stage-duration distributions (de Valpine et al. 2014) may be specified by either specifying the stage-duration distribution itself, *f*(*t*), or by specifying the corresponding hazard function *h*(*t*) (see relationships in section 1.4.1.5). If individuals can leave a certain state to *K* > 1 other states (‘death’ being one of them), the unconditional state-duration distribution is the probability distribution corresponding to the total hazard rate 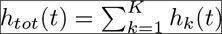, where *h*_*k*_(*t*) are the transition specific hazard rates. It is, however, often of greater interest to study the probability distributions for age or time of state-transition conditional on survival. For example, Ergon et al. (2009) estimated the latent distributions [^iv^ By ‘latent distributions’ we here mean the distribution of the life-history trait that is independent of mortality, and this distribution represent the trait that is under selection. In contrast, the ‘realized distribution’ is the distribution of time/age of maturation among those individuals that survive until maturation. Because individuals that have a late latent maturation need to survive over a longer period before maturation can be observed, these individuals will be under-represented in the realized distribution (unless individual time/age of maturation is strongly negatively correlated with mortality hazard rate).] for seasonal timing of onset of reproduction in populations of overwintering field voles.

Matrix population models are memory-less (Markovian) in the sense that state transition probabilities only depend on the current state of the individual. However, in individual-based “i-state configuration models” (Caswell & John 1992) one can track individual time since entering specific states, and hazard rates may in addition be dependent on “global time” and age of each individual. Such models could potentially be used to compute full conditional likelihoods (Carlin & Louis 2009), and modell fitting of capture-recapture data could be done by using Bayesian MCMC sampling. Potential applications could include estimation of (mortality) cost or reproduction depending on time since conception (or maturation), or mortality rates depending on time since infection of a pathogen.

Whenever individuals can transition from one state to more than one other state (including mortality), there is a competing risk situation among the potential transitions (Appendix S2). In addition, individuals may transition from one state to another state (or back to the original state) via other states during a given time period. A general, and very useful, tool to compute transition probability matrices from transition hazards between any number of states, involve the use of matrices of transition hazards and matrix exponentials (Keyfitz & Caswell 2005 chap. 17.5; Miller & Andersen 2008; Conn et al. 2012): First, a matrix **A** is constructed with transition hazards as the off-diagonal elements, and the negative sum of all transition hazards out of the given state as the diagonal elements, such that each row in **A** sums to zero. The corresponding transition probability matrix for an interval of length Δ is then simply

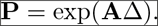

exp refers here to a matrix exponential; function “expm” in the “expm” package of R (Goulet et al. 2017) and function “MatrixExponential” in the LinearAlgebra package in Maple™. See Miller & Andersen (2008) and Conn et al. (2012) for various examples. To include continuous hazard functions of time, or time since entering the states (as discussed above), one may discretize by computing transition probability matrices over many short time-intervals (e.g. days), assuming constant hazard rates within each interval.

## 1.4.2 Figures 5 and 6

Figures 5 and 6 illustrate the differences between elasticities of λ to *S* and to *h̄* by using a very simple two-stage population model without density dependence. With a post-reproductive census (Caswell 2001, p. 27), this model can be written as

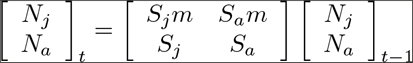

where indices *j* and *a* represent respectively ‘juveniles’ (newborn) and ‘adults’ (1 year or older), *N* is number of individuals, *S* is survival probability and *m* is fecundity. Index value *t* is the time-step (year).

With a pre-reproductive census, there are no counts of newborn, and the dynamics of the population is simply described by

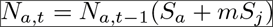

Hence, the population growth rate in this model is

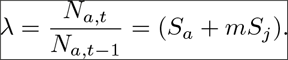

From this we can calculate elasticities of λ to survival probabilities and their corresponding time averaged mortality hazard-rates,

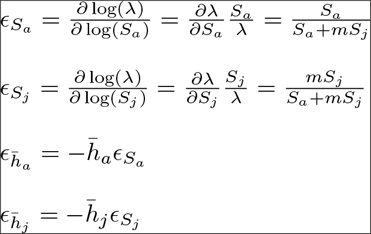

The elasticity ratio with respect to survival probabilities is thus

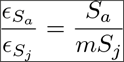

(i.e., *ϵ*_*S*_*a*__ is grater that *ϵ*_*S*_*j*__ when *S*_*a*_ > *mS*_*j*_). The elasticity ratio with respect to time-averaged mortality hazard rates is

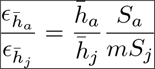

Figure 5 in the main text shows the effect of reparameterizing the model with survival probabilities representing different interval lengths, and Figure 6 compares elasticity ratios with respect to survival probabilities and time-averaged mortality hazard rates. Although these figures could have been produced simply by using the relationships derived for the two-stage model above, we have chosen to do it in a more general way using the ‘vitalsens’ function in the ‘popbio’ R-package (Stubben & Milligan 2007). Hence, the code below can quite easily be adapted to any matrix population model or to other kinds or reparameterizations.

## 1.4.2.1 Code - Figure 5

~~~
library(popbio) *# For function ‘vitalsens’*

*# Function for elastisities with reparameterization of survival probabilities*
*# representing different interval lengths*
ela_S = function(
    Sj.y, *# Yearly juveile survival*
    Sa.y, *# Yealry adult survival*
    m, *# Fecundity*
    t *# No. time units per year*
){
 el = expression(*# Expressions for each of the matrix elements in a vector (by row)*
   Sj.t^t*m, Sa.t^t*m,
   Sj.t^t, Sa.t^t
)
 vr = list(Sj.t=Sj.y^(1/t), Sa.t=Sa.y^(1/t), m=m, t=t) # Num. values for demogr. params.
 v = vitalsens(el, vr)
 elas = v$elasticity
 names(elas) = rownames(v)
 return(elas)
}

*# Above function allowing vector inputs for t*
Ela_S = Vectorize(ela_S, “t”)

*# Using same structure to also reparameterize survival probabilities with respect to*
*# time averaged hazard rates*
*# This can be substantially simplified, but keeping the general structure as an example*
*# and for validation*
ela_h = function(Sj.y, Sa.y, m, t){
  el = expression(
   exp(-hj.t)^t*m, exp(-ha.t)^t*m,
   exp(-hj.t)^t, exp(-ha.t)^t
 )
  vr = list(hj.t=-log(Sj.y^(1/t)), ha.t=-log(Sa.y^(1/t)), m=m, t=t)
  v = vitalsens(el, vr)
  elas = v$elasticity
  names(elas) = rownames(v)
  return(elas)
}
*# Above function allowing vector inputs for t*
Ela_h = Vectorize(ela_h, “t”)

*# Function for plotting*
plot.panel = function(
    Sj.y, *# Juvenile yearly survival probability*
    Sa.y, *# Adult yearly survival probability*
    m, *# Fecundity*
    il *# Vector of interval lengths in unit of days for reparameterization*
 ){
  ES = Ela_S(Sj.y, Sa.y, m, 1/il)
  Eh = Ela_h(Sj.y, Sa.y, m, 1/il)
  plot(1,1, type=“n”, ylim=c(0,3), xlim=range(il), axes=F,
       xlab = “Survival interval length”,
       ylab = expression(paste(“Elasticity of”, lambda)))
  box()
  axis(2)
  axis(1, at = c(7/365, 1/4, 1/2, 1, 2), labels=c(“1w”, “3m”, “6m”, “1y”, “2y”))
  abline(v = c(7/365, 1/4, 1/2, 1, 2), col=“grey”)
  lines(il, ES[“Sj.t”,], col=“blue”, lty=2)
  lines(il, ES[“Sa.t”,], col=“red”, lty=2)
  lines(il, ES[“m”,], col=“green”)
  lines(il, −Eh[“hj.t”,], col=“blue”)
  lines(il, −Eh[“ha.t”,], col=“red”)
  legend(“topright”, title = “ Elasticity to …”, title.adj=0, bg=“white”,
         legend=c(“Juvenile survival”, “Adult survival”, “Juvenile mortality”,
“Adult mortality”, “Fecundity”),
         col=c(“blue”,“red”,“blue”,“red”,“green”), lty=c(2,2,1,1,1))
  title(paste(“Juvenile survival = ”, Sj.y, “\nAdult survival = ”, Sa.y, sep=“”))
}

*# Plotting:*
par(mfrow = c(1,2))
plot.panel(Sj.y = 0.2, Sa.y = 0.6, m = 2, il = (1:(2*385))/365)
title(“(a)”, adj=0)
plot.panel(Sj.y = 0.6, Sa.y = 0.2, m = 2, il = (1:(2*385))/365)
title(“(b)”, adj=0)
~~~

## 1.4.2.2 Code - Figure 6

~~~
*# Using functions from Fig. 5*

library(RColorBrewer) *# for brewer.pal()*

*# Creating martices with elasticity ratios*
n = 100
m = 2
Sj = seq(0.01,0.99,length.out=n)
Sa = seq(0.01,0.99,length.out=n)
Er.S = Er.h = matrix(NA, n, n)
for(i in 1:n){
  for(j in 1:n){
    ES = ela_S(Sj[i], Sa[j], m, 1)
    Eh = ela_h(Sj[i], Sa[j], m, 1)
    Er.S[i,j] = ES[“Sa.t”]/ES[“Sj.t”]
    Er.h[i,j] = Eh[“ha.t”]/Eh[“hj.t”]
  }
}

*# Plotting*
par(mfrow=c(1,2)) *# Does not work with filled.contour()*
Col = c(rev(brewer.pal(9,“Blues”)), brewer.pal(9,“Reds”))
maxval = 6
bins = seq(−maxval, maxval, length.out=19)

filled.contour(Sj, Sa, log(Er.S), levels=bins, nlevels=18, col=Col,
        plot.axes={axis(1); axis(2); text(c(0.2, 0.6), c(0.6, 0.2), c(“a”,“b”),
cex=1.3)},
        plot.title = title(“Elasticity ratio - survival probability”, cex.main=1.1,
xlab=“Juvenile survival probability”, ylab=“Adult survival probability”),
        key.title = title(main = “ln(ratio)”, cex.main=0.9, line=1))
title(“(a)”, outer=T, adj=0.05, line=−2)
filled.contour(Sj, Sa, log(Er.h), levels=bins, nlevels=18, col=Col,
        plot.axes={axis(1); axis(2); text(c(0.2, 0.6), c(0.6, 0.2), c(“a”,“b”), cex=1.3)},
        plot.title = title(“Elasticity ratio - - mortality rate”, cex.main=1.1,
                                              xlab=“Juvenile survival probability”, ylab=“Adult survival probability”),
        key.title = title(main = “ln(ratio)”, cex.main=0.9, line=1))
title(“(b)”, outer=T, adj=0.05, line=−2)
~~~

## 1.4.3 Figure 7

As shown in Figure 6 above, patterns in elasticities to mortality hazard rate are very different from patterns in elasticities to survival probability. One way to understand this is to note, as shown in Figure 7, that the relationship between log(*S*) and log(*h*) is very non-linear (log(*S*) = — exp(log(*h*)) (the first derivative in this plot is — *h*). When the hazard rate is low (survival is high), a large relative change in the hazard rate (to the left in the figure) corresponds to a small relative change in survival probability. When the hazard rate is high (survival is low), a small relative change in the hazard rate corresponds to a large relative change in survival. This is also reflected in the relationship between elasticities to survival probability and elasticity to mortality hazard rate given above; *ϵ*_*h̄*_ = —*hϵ*_*S*_.

## 1.4.3.1 Code - Figure 7

~~~
S = seq(0.001, 0.999, length.out=40)
y = log(S)
x = log(−log(S))

par(mai=c(1.2,1.2,1.2,1.2))
plot(x,y, xlab=“ln(Mortality hazard rate)”, ylab=“ln(Survival probability)”, type=“n”)

lS = c(0.001, 0.01, 0.1, 0.5, 0.9)

axis(3, at = log(−log(lS)), labels=as.character(lS), col.ticks=“darkgrey”,
     col.axis=“darkgrey”)
axis(4, at = log(lS), labels=as.character(lS), col.ticks=“darkgrey”, col.axis=“darkgrey”)
abline(h=log(lS), v=log(−log(lS)), col=“darkgrey”)
mtext(“Survival probability”,3, line=2.5, col=“darkgrey”)
mtext(“Survival probability”,4, line=2.5, col=“darkgrey”)
lines(x,y, lwd=2)
~~~

## 1.4.4 Variance-stabilized sensitivities

In the final paragraph of this section in the main text, we comment on the use of variance-stabilized sensitivities (VSS) as defined by Link & Doherty (2002). VSS are based on an assumed relationship between the temporal variance of a demographic parameter *θ* and the temporal mean *μ* of this parameter, which may be described as Var(θ) = *f*(*μ*). The delta-method approximation to the variance of a transformation *q*(*θ*) of this parameter is then

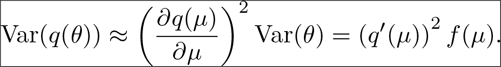

For a given variance-mean relationship *f*(*μ*) the transformation *q*(*θ*) is then said to be a variance-stabilizing transformation when Var(*q*(*θ*)) is independent of *μ*, i.e., when (*q*′(*μ*))^2^ *f* (*μ*) is a constant. Link & Doherty (2002) then defined the variance-stabilized sensitivity for a given variance-mean relationship as the sensitivity of log(λ) to the transformed parameter *q*(*θ*)

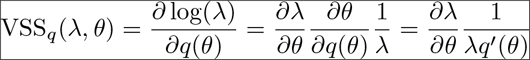

To find the variance-mean relationship that corresponds with a particular variance stabilizing transformation *q*(*θ*), we need to solve (*q*′(*μ*))^2^ *f* (*μ*) = *C* for *f*(*μ*) where *C* is an arbitrary positive constant. When *q*(*θ*) is a logit-transformation of survival probability, we have 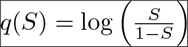 and hence

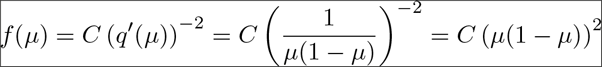

Similarly, the loglog-link on survival probability, *q*(*S*) = log(− log(*S*)), is a variance stabilizing transformation when

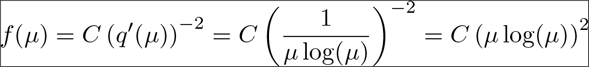

When using the loglog-link on survival probability, *q*(*S*) = log(− log(*S*)) = log(*h̄*), as a variance stabilizing transformation, the variance-stabilized sensitivity becomes

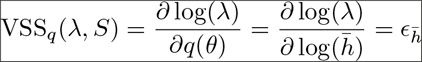

Hence, using the loglog-link to obtain a variance-stabilized sensitivity to survival probability, assuming that the temporal variance in interval specific survival probabilities is proportional to (*μ*log(*μ*))^2^, is equivalent to using the elasticity to time-averaged mortality hazard rate.

## Appendix S2: Competing risks and cause-specific mortality probabilities

**Figure S2.1:**
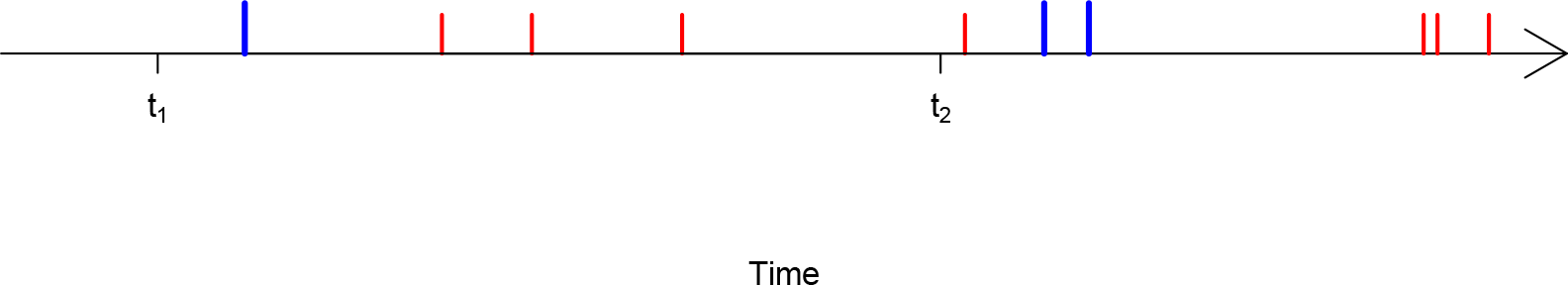
Realized times of two types of independent recurrent events (or “deadly events experienced by an immortal individual“)

Figure S2.1 illustrates a situation where another (“blue”) cause of mortality has been added to the “red” cause of mortality illustrated in the figure in Box 1 in the main text. If we, for simplicity, assume that the hazard rates for these two causes of mortality are constant over an interval with values *h*_1_ and *h*_2_, the total mortality hazard rate is now *h*_1_ + *h*_2_, and the probability of surviving the interval of length Δ is 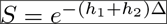.

It is essential to realize that adding the blue cause of mortality in this example reduces the probability of dying from the red cause, even though the hazard rate representing the red cause remains the same. This is due to the fact that the individual now may die from a blue cause before a red event takes place. The probability of dying from a specific cause (often referred to as a “risk”), when the hazard rate of this cause remains unchanged, will always go down when the hazard rate of other mortality causes increase (or go up if the hazard rate of other causes decrease). One may say that the risks of death are “competing”, and the probability of “winning” depends on the strength of the competitors (i.e., mortality probabilities are intrinsically linked). Hence, a change in the probability of dying of a specific cause may be due to changes in any of the cause-specific mortality hazard rates. When studying e.g., environmental influence on different causes of mortality, or how different causes of mortality vary and co-vary among groups of individuals or among years within a population, it is therefore of prime interest to estimate the hazard rates of different causes of mortality rather than the probabilities of dying from different causes.

Discrete-time models (such as classical capture recapture models) that incorporate different causes of mortality involve state-transition probability parameters representing the probabilities that an individual that is alive at one time-point (*t*_1_) is either alive or dead from specific causes at a later point in time (*t*_2_). These transition probabilities should sum to 1. Hence, 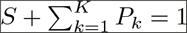, where *S* is the survival probability and the *P*_*k*_’s are the probabilities of dying from different causes classified in *K* categories. Assuming, for simplicity, that each of the cause-specific hazard rates, *h*_*k*_ are constant within intervals, we know from above that 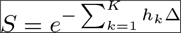. To find the probability of dying from cause *k*, *P*_*k*_, we may use the definition of hazard rates as

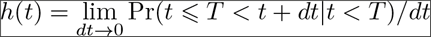

(Collett 2014) where *T* is time of death and *dt* is an infinitesimally small unit of time (i.e., *h*(*t*)*dt* is the probability that the individual will die before time *t* + *dt* given that it is alive at time *t*). From this we see that the relationship between the hazard rate and the probability density for time of death at time *t*, *f*(*t*), is

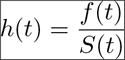

where *S*(*t*) is the probability of being alive at time *t*, which equals 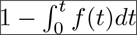. Using this relationship, we can now find the probability density function for the time of death from cause *k*, *f*_*k*_(*τ*), where *τ* = *t* — *t*_1_ is the time since the beginning of the interval,

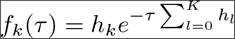

Note that this density function is not a proper probability distributions function as it does not integrate to 1 over infinite time. Instead, it integrates to the probability of ever dying from cause *k*, which in the case of constant hazard rates becomes 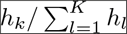. To find the probability of dying from cause *k* before the end of the interval, we need to integrate *f*_*k*_(*τ*) from *τ* = 0 to *τ* = Δ (= *t*_2_ — *t*_1_), and get

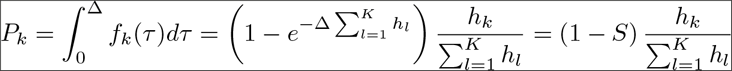

This expression can be generalized to cases where cause-specific hazard rates may vary over time but remain proportional throughout the interval such that 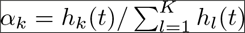, in which case we get

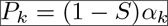

Note that the conditional probability of having died from cause *k*, given that the individual has died, in the case of proportional hazard rates is the fraction of the total mortality hazard rate attributed to the specific cause, *α*_*k*_, which in this case equals the probability of ever dying of this cause. This is, however, not the case when hazard rates are not proportional over time.

Section 1.4.1.6 in Appendix S1 outlines a more general approach to deriving/computing transition probabilities under competing risk.

## Appendix S3: Fitting capture-recapture models with cause-specific mortality

## 3.1 Introduction

When information about cause of death (e.g., harvesting) is available for at least a subset of marked individuals in the population, multi-state capture-recapture models can be fitted to the data to address questions relating to e.g. compensatory effects of harvesting or age-related patterns in mortality from different causes. However, commonly used software for fitting multi-state capture-recapture data such as MARK (White & Burnham 1999) and E-SURGE (Choquet, Rouan & Pradel 2009) use the multinomial logit-link function to constrain rows of the transition matrix to sum to 1. For example, the probability of transitioning from the alive state to death-by-cause state *k* (i.e., probability of dying from cause *k*) may be modelled as

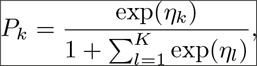

while the probability of surviving is

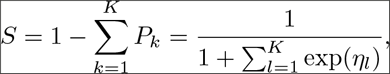

where *η*_*k*_ is the linear predictor relating to cause *k*. This multinomial link-function does not facilitate modelling of cause-specific hazard rates (Appendix S2). Instead, based on the expressions for *S* and *P*_*k*_ derived in Appendix S2, we can model log-linear effects on the mortality hazard rates by using the multinomial link-function defined by

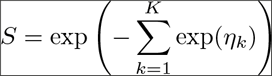

and

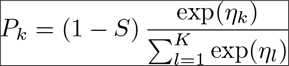

To fit, for example, a model with two causes of mortality one needs a 4 by 4 transition matrix

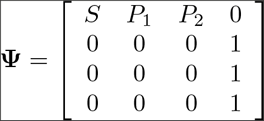

The four states are here ‘alive’, ‘newly dead from cause 1’, ‘newly dead from cause 2’ and a final absorbing state (rows represent “from-state” and columns represent “to-state”). The distinction between newly dead states and the final state is necessary because individuals must be reported dead in the year they died (i.e., year of death of recovered individuals must be known). Recapture/recovery probabilities of the four states are

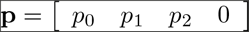

In the most general (full, unconstrained) model, parameters would be individual and year specific, but indices are omitted here for simplicity. Survival probabilities in the general multi-state capture-recapture models must be fixed to 1 for the three first states and to 0 (or any probability value) for the absorbing state.

To facilitate modelling of mortality hazard rates, we modified the functions used for fitting multi-state capture-recapture models in the R-package ‘marked’ package (Laake, Johnson & Conn 2013). This package combines a system for highly flexible model formulations (including modelling of covariate effects relating to time, age, cohort and individuals), similar to the popular ‘RMark’ package, with highly efficient numerical optimization of the likelihood function using automatic differentiation through executables compiled by the ADMB software (Fournier et al. 2012). The files are available on https://github.com/torbjore/study_mortality_with_hazard_rates, and a worked example using simulated data is included below.

Schaub & Pradel (2004) and Schaub (2009) noted that only some sub-models of another version of the general multi-state capture-recapture model (see Table 1 in the main text) are identifiable. This is also the case for our model, but a detailed treatment of identifiability is beyond our scope here.

## 3.2 Preparation

To use the modified ‘marked’ files, an executable must first be compiled and saved on your computer. To do this, first install the ADMB software from http://www.admb-project.org/. ADMB needs a C++ compiler, but this is included in some of the ADMB installation files. It is safest to install ADMB in a directory without spaces in its name and path. The code below assumes that ADMB is installed in “C:/admb”, but this can be modified. Once ADMB and a C++ compiler is installed, the following code will create a ‘multistate_csmhr.exe’ file in your working directory (use install.packages() to install any missing packages).

~~~
library(marked)
library(R2admb)
library(RCurl) # *For getURL*

prepare_admb = function()
{
  Sys.setenv(PATH = paste(“c:/admb/bin;c:admb/utilities;c:/admb/utilities/mingw64/bin;”,
  Sys.getenv(“PATH”), sep = “;”))
  Sys.setenv(ADMB_HOME = “c:/admb”)
  invisible()
}
prepare_admb()

*# Downloading the ADMB tpl file and building a local executable*
*# Breaking up string to fit in box*
reprstr1 = “https://raw.githubusercontent.com/torbjore/study_mortality_with_hazard_rates/”
reprstr2 = “master/modified_marked_files/”
tplfile = paste(reprstr1, reprstr2, “multistate_csmhr.tpl”, sep=“”)
tpl = getURL(tplfile)
write(tpl, “multistate_csmhr.tpl”)
marked::setup_admb(“multistate_csmhr”, compile=TRUE, clean=FALSE)
~~~

## 3.3 Example

In this simple example we will simulate 10 years of capture-recapture data with two causes of mortality; ‘hunting’ and ‘natural’. We will assume that both mortality hazard rates varies independently over time, that probability of recapture of an individual alive is 0.5, that the reporting probability of a hunted individual is 0.7, and that the probability that an individual which died of other (natural) causes will be found and reported is 0.1.

First, we simulate data:

~~~
set.seed(11) *# for reproducibility*

p = c(0.5, 0.7, 0.1, 0)
N.ind = 200 *# Number of individuals marked each year*
N.yr = 10 *# Number of years*
*# Year-specific hazard rates for the two causes*
h1 = runif(N.yr-1, 1/25, 1/2)
h2 = runif(N.yr-1, 1/25, 1/2)

*# For computation of transition matrix*
fTM = function(h1, h2){
  *# Yearly survival probability*
  S = exp(−(h1+h2))
  *# Yearly mortality probabilities*
  P1 = (1-S)*h1/(h1+h2)
  P2 = (1-S)*h2/(h1+h2)
  *# Transition matrix*
  tm = matrix(
  c(
   S, P1, P2, 0,
   0, 0, 0, 1,
   0, 0, 0, 1,
   0, 0, 0, 1
 ), 4, 4, byrow = T
)
 return(tm)
}

first = matrix(rep(1:(N.yr - 1), each = N.ind), ncol=1) *# When marked*

*# Simulating Matrix of true states*
N.ind.tot = length(first)
St = matrix(NA, N.ind.tot, N.yr)
for(i in 1:N.ind.tot){
  St[i,first[i]] = 1 *# All individuals introduced as alive*
  for(j in first[i]:(N.yr-1)){
    St[i,j+1] = sample(1:4, 1, prob = fTM(h1[j], h2[j])[St[i,j],])
  }
}
St[is.na(St)] = 4 *# to get zero capture probability before marking*

*# Computing matrix of state-dependent capture probabilities*
P = matrix(p[St], N.ind.tot, N.yr)

*# Simulating Capture History data*
capt = rbinom(N.ind.tot*N.yr,1,P)
CH = St*capt
for(i in 1:N.ind.tot){
  CH[i,first[i]] = 1 *# Conditioning on first capture; hence inds are always seen when marked*
}
~~~

Then we source in the modified marked functions and prepare the data for model fitting:

~~~
library(marked)
library(R2admb)
prepare_admb() *# Function defined above*

*# Source in modified functions from Git-hub*
crm_csmhr_FUN = paste(reprstr1, reprstr2, “crm_csmhr.R”, sep=“”)
mscjs_csmhr_FUN = paste(reprstr1, reprstr2, “mscjs_csmhr.R”, sep=“”)
source(crm_csmhr_FUN)
source(mscjs_csmhr_FUN)

*# Transforming data to a sting variable for ‘marked’*
simdata = data.frame(
  ch = apply(CH, 1, function(i) paste(c(“0”,“A”,“B”,“C”)[i+1], collapse = “”))
)
simdata$ch = as.character(simdata$ch)

*# Adding an individual seen in state D at the *last* occasion to get the right dimension*
*# of the transitioin matrix. Since this individual is captured only in the last year,*
*# and we condition on first capture, this individual will not affect estimates (and will*
*# be removed internally)*

simdata = rbind(simdata, ch = paste(paste(rep(“0”, N.yr-1), collapse = “”), “D”, sep=“”))

simdata.processed = process.data(simdata,model=“Mscjs”,strata.labels=c(“A”,“B”,“C”,“D”))
simdata.ddl = make.design.data(simdata.processed)
~~~

The resultant `simdata.dll` object is a list of data frames with predictor variables for all parameters in the full (unconstrained) model; one data.frame for modelling of log hazard rates (`simdata.ddl$Psi`), and one data.frame for modelling of logit recapture/recovery probabilities (`simdata.ddl$p`). Individual covariates, grouping variables and individual ages at first capture may be included in `process.data()` (see `help(package=“marked”`) for documentation).

Below, we create variables `simdata.ddl$Psi$hunting` and `simdata.ddl$Psi$natural`, use this to constrain the model and fit it by using the modified functions:

~~~
simdata.ddl$Psi$hunting = ifelse(simdata.ddl$Psi$tostratum == “B”, 1, 0)
simdata.ddl$Psi$natural = ifelse(simdata.ddl$Psi$tostratum == “C”, 1, 0)

*# Submodel constraints*
model = list(
  p = list(formula = ~ −1 + stratum),
  Psi = list(formula = ~ −1 + natural:time + hunting:time)
)

*# NB! Convergence may be sensitive to starting values. It’s a good idea to try many*
*# different starting values to check for convergence at local maxima:*
inits = list(p=rep(0,3), Psi = rep(mean(log(c(h1,h2))), 2*(N.yr-1)))

mod1_csmhr = crm_csmhr(simdata.processed, simdata.ddl, clean=F, model.parameters=model,
initial=inits, hessian=TRUE)

##
## Elapsed time in minutes: 0.1247
*# NB! Run with clean=F, or else the executable will be deleted!*
~~~

Note that is not necessary to fix the survival probabilities or fixed transition matrix elements as this is done internally in the `crm_csmhr` function.

To see parameter estimates and a summary of the model fit, type the name of the model object

~~~
mod1_csmhr
##
## crm Model Summary
##
## Npar: 21
## −2lnL: 8687.64
## AIC: 8729.64
##
## Beta
## Estimate se lcl ucl
## p.stratumA −0.04400161 0.05217081 −0.1462564 0.05825318
## p.stratumB 0.54749248 0.39423422 −0.2252066 1.32019156
## p.stratumC −2.21528017 0.17240226 −2.5531886 −1.87737173
## Psi.natural:time1 −2.13366000 0.53993877 −3.1919400 −1.07538000
## Psi.natural:time2 −2.54077677 0.38698251 −3.2992625 −1.78229105
## Psi.natural:time3 −1.76863318 0.29324920 −2.3434016 −1.19386475
## Psi.natural:time4 −0.63110307 0.10472320 −0.8363605 −0.42584559
## Psi.natural:time5 −0.97332549 0.13576997 −1.2394346 −0.70721634
## Psi.natural:time6 −1.16448218 0.25093775 −1.6563202 −0.67264419
## Psi.natural:time7 −1.25585599 0.17927273 −1.6072305 −0.90448144
## Psi.natural:time8 −1.38657207 0.22280072 −1.8232615 −0.94988265
## Psi.natural:time9 −2.02119027 0.47154286 −2.9454143 −1.09696626
## Psi.time1:hunting −1.32971130 0.24566097 −1.8112068 −0.84821580
## Psi.time2:hunting −2.93815957 0.32592799 −3.5769784 −2.29934071
## Psi.time3:hunting −1.20405839 0.17919551 −1.5552816 −0.85283519
## Psi.time4:hunting −2.85968339 0.30819935 −3.4637541 −2.25561266
## Psi.time5:hunting −2.49942724 0.26982471 −3.0282837 −1.97057081
## Psi.time6:hunting −0.77625675 0.17492928 −1.1191181 −0.43339537
## Psi.time7:hunting −2.15652028 0.24773121 −2.6420735 −1.67096710
## Psi.time8:hunting −1.65644715 0.20087128 −2.0501549 −1.26273944
## Psi.time9:hunting −0.66242305 0.17041356 −0.9964336 −0.32841248
~~~

Here, `p` parameters are logit-linear coefficients in the model for recapture/recovery probabilities, and `Psi` parameters are coefficients in the log-linear models for each of the mortality hazard rates.

Please note that convergence on a local likelihood peak is not unlikely, and it is highly recommendable to try many different starting values (lower AIC values indicate higher likelihood values). It is also possible to change convergence criteria and perform various diagnostics (as well as computing likelihood profiles, confidence intervals by MCMC, etc) by passing ADMB command line options through the extra.args argument in `crm_csmhr` or by modifying the `multistate_csmhr.tpl` file - see the ADMB manual at http://www.admb-project.org/.

The `marked::predict.crm` function cannot be used for prediction of mortality hazard rates (only for recapture/recovery probabilities), but we include a simple function for prediction below. Here we compute and plot the predicted mortality hazard rates and compare them to the true values used in the simulation:

~~~
predict.marked.csmhr = function(fit, newdata, par, invlink){
  model = fit$model.parameters[[par]]$formula
  X = model.matrix(model, newdata)
  b = fit$results$beta[[par]]
  vc = fit$results$beta.vcv
  nms = dimnames(vc)[[1]]
  use = substring(nms,1,nchar(par)) == par
  vc = vc[use,use]
  eta = X %*% b
  se = sqrt(diag(X %*% vc %*% t(X)))
  D = newdata
  D$eta = eta
  D$se = se
  D$est = invlink(eta)
  D$lwr = invlink(eta - 1.96*se)
  D$upr = invlink(eta + 1.96*se)
  return(D)
}

D = expand.grid(year = 1:9, cause = c(“hunting”,“natural”))
D$time = factor(paste(“time”, D$year, sep=“”))
D$hunting = ifelse(D$cause == “hunting”, 1, 0)
D$natural = ifelse(D$cause == “natural”, 1, 0)

Pred = predict.marked.csmhr(fit = mod1_csmhr, newdata = D, par = “Psi”,
                                   invlink = function(x) exp(x))

*# Adding true values*
Pred$true = ifelse(Pred$hunting==1, h1, h2)

*# Plotting*
library(plotrix) *# for plotCI*

par(mfrow=c(1,2))
Ylim = range(c(Pred$lwr, Pred$upr))
with(Pred[Pred$cause==“hunting”,],
     plotCI(year, est, ui=upr, li=lwr, gap=T, ylim=Ylim,
            xlab = “Year”, ylab = “Mortality hazard rate”))
with(Pred[Pred$cause==“hunting”,], lines(year, est, type=“b”))
with(Pred[Pred$cause==“hunting”,], points(year,true, pch=16, col=“red”))
title(“Hunting mortality”)

with(Pred[Pred$cause==“natural”,],
     plotCI(year, est, ui=upr, li=lwr, gap=T, ylim=Ylim,
            xlab = “Year”, ylab = “Mortality hazard rate”))
with(Pred[Pred$cause==“natural”,], lines(year, est, type=“b”))
with(Pred[Pred$cause==“natural”,], points(year,true, pch=16, col=“red”))
title(“Natural mortality”)
~~~

We do the same for recapture/recovery probabilities:

~~~
*# Recapture/recovery probability*
D.p = data.frame(stratum = c(“A”,“B”,“C”))
Pred.p = predict.marked.csmhr(fit = mod1_csmhr, newdata = D.p, par = “p”,
                                     invlink = function(x) exp(x)/(1+exp(x)))
Pred.p$true = p[1:3]

Ylim = c(0,1)
with(Pred.p,
     plot(1:3, est, ylim=Ylim, xlim=c(0.7,3.3),
          xlab = “”, ylab = “Recapture/recovery probability”, type=“n”, axes=F))
with(Pred.p, plotCI(1:3, est, add=T, ui=upr, li=lwr, gap=T))
with(Pred.p, points(1:3, true, pch=16, col=“red”))
box()
axis(2)
axis(1, at=1:3, labels = c(“Alive”, “Hunted”, “Naturally dead”))
~~~

Finally, we use the delta-method to compute approximate standard errors of the log of total mortality hazard rates and plot the confidence intervals of survival probabilities. At the same time, we compute and plot estimates of the sampling correlation between the two log hazard rates for each year (may indicate identifiability problems if highly negative).

~~~
*# Function for a single year*
S.delta = function(yr, fit, D){
  DD = D[D$year==yr,]
  model = fit$model.parameters[[“Psi”]]$formula
  X = model.matrix(model, DD)
  b = fit$results$beta[[“Psi”]]
  vc = fit$results$beta.vcv
  nms = dimnames(vc)[[1]]
  use = substring(nms,1,3) == “Psi”
  vc = vc[use,use]
  log.h = X %*% b
  VC.log.h = X %*% vc %*% t(X)
  h = exp(log.h)
  h.tot = sum(h)
  log.h.tot = log(h.tot)
  x = h/h.tot
  var.log.h.tot = t(x) %*% VC.log.h %*% x
  se.log.h.tot = sqrt(diag(var.log.h.tot))
  lwr.log.h.tot = log.h.tot - 2*se.log.h.tot
  upr.log.h.tot = log.h.tot + 2*se.log.h.tot
  list(
    S = exp(−h.tot),
    lwr.S = exp(−exp(upr.log.h.tot)),
    upr.S = exp(−exp(lwr.log.h.tot)),
    cor.log.h = cov2cor(VC.log.h)[1,2]
 )
}

S.est = sapply(1:(N.yr-1), S.delta, fit=mod1_csmhr, D=D)
S.est = as.data.frame(t(S.est))

par(mfrow=c(1,2))
plot(1:(N.yr-1), unlist(S.est$S), ylim=c(0,1),
     xlab = “Year”, ylab = “Survival probability”)
with(S.est,
     plotCI(1:(N.yr-1), unlist(S), ui=unlist(upr.S), li=unlist(lwr.S), add=T, gap=T))
lines(1:(N.yr-1), unlist(S.est$S), type=“b”)
points(1:(N.yr-1),exp(−(h1+h2)), pch=16, col=“red”)
title(“Survival probability”)

plot(1:(N.yr-1), unlist(S.est$cor.log.h), ylim=c(−1,1),
     xlab = “Year”, ylab = “Correlation”, type=“b”)
title(“Sampling correlation\n between log hazard estimates”)
~~~

